# MR-based camera-less eye tracking using deep neural networks

**DOI:** 10.1101/2020.11.30.401323

**Authors:** Markus Frey, Matthias Nau, Christian F. Doeller

## Abstract

Viewing behavior provides a window into many central aspects of human cognition and health, and it is an important variable of interest or confound in many fMRI studies. To make eye tracking freely and widely available for MRI research, we developed DeepMReye: a convolutional neural network that decodes gaze position from the MR-signal of the eyeballs. It performs camera-less eye tracking at sub-imaging temporal resolution in held-out participants with little training data and across a broad range of scanning protocols. Critically, it works even in existing datasets and when the eyes are closed. Decoded eye movements explain network-wide brain activity also in regions not associated with oculomotor function. This work emphasizes the importance of eye tracking for the interpretation of fMRI results and provides an open-source software solution that is widely applicable in research and clinical settings.

## Introduction

Eye movements are a direct expression of our thoughts, goals and memories and where we look determines fundamentally what we know about the visual world. The combination of eye tracking and neuroimaging can thus provide a window into many central aspects of human cognition, along with insights into neurodegenerative diseases and neural disorders of the brain (Anderson & MacAskill, 2013). A widely used tool to study human brain function is functional magnetic resonance imaging (fMRI), which allows examining brain activity while participants engage in a broad range of tasks. Viewing behavior is either a variable of interest or one of potential confound in many fMRI studies, yet the very large majority of them does not perform eye tracking.

We argue that eye tracking can and should be a central component of fMRI research. Not only does it allow in-depth insights into brain function, but it also offers a powerful behavioral read-out during scanning. Importantly, eye movements are also associated with perceptual distortions (Morrone et al., 2005), visual and motor activity (Berman et al., 1999; Petit & Haxby, 1999) and imaging artifacts (McNabb et al., 2020), which can severely affect the interpretation of neuroimaging results. If differences in viewing behavior between experimental conditions remain undetected, there is a high risk of misinterpreting differences in the observed brain activity. Crucially, this is not restricted to studies of the visual system but affects task-based and resting-state neuroimaging on a large scale.

One example that illustrates the importance of eye tracking also for studies of higher-level cognition is the subsequent-memory effect, the observation that hippocampal activity during encoding reflects whether a stimulus is later remembered or forgotten (Wagner et al., 1998). This effect is often attributed to mnemonic processes in the hippocampus. However, because we also tend to remember images better that we visually explored more thoroughly (Kafkas & Montaldi, 2011) and because hippocampal activity scales with the number of fixations on an image (Liu et al., 2017), the interpretation of hippocampal activity in this context can be difficult. In many such cases, it remains unclear if the observed brain activity reflects higher-level cognitive operations or if it is driven by viewing behavior (Voss et al., 2017).

MR-compatible camera eye trackers offer a solution. They track gaze position during scanning and hence allow to analyze or account for gaze-related brain activity. In practice, however, camerasystems are not applicable in many research and clinical settings, often because they are expensive, require trained staff and valuable setup and calibration time, and because they impose experimental constraints (e.g. the eyes need to be open). Moreover, they cannot be used in visually impaired patient groups or post-hoc once the fMRI data has been acquired.

An alternative framework is MR-based eye tracking: the reconstruction of gaze position directly from the MR-signal of the eyeballs. While previous work suggested that this is indeed feasible (Tregellas et al., 2002; Beauchamp, 2003; Heberlein et al., 2006; Son et al., 2020), several critical constraints remained that limited the usability to specific scenarios. These earlier approaches were not as accurate as required for many studies, were limited to the temporal resolution of the imaging protocol, and most importantly required dedicated calibration scans for every single participant.

Here, we present DeepMReye, a novel open source camera-less eye tracking framework based on a convolutional neural network (CNN) that reconstructs viewing behavior directly from the MR-signal of the eyeballs. It can be used to perform highly robust camera-less eye tracking in future fMRIexperiments, but importantly also in datasets that have already been acquired. It decodes gaze position in held-out participants at sub-imaging temporal resolution, performs unsupervised outlier detection and is robust across a wide range of viewing behaviors and fMRI protocols. Moreover, it can create new experimental opportunities for example by performing eye tracking while the eyes are closed (e.g. during resting-state or REM-sleep) or in patient groups for which eye-tracker calibration remains challenging.

## Results

In the following, we present our model and results in three sections. First, we introduce our datasets, tasks, data processing pipeline and CNN in detail. Second, we show that the decoded gaze positions are highly accurate and explore the applicability and requirements of DeepMReye in depth. Lastly, by regressing the decoded gaze labels against the simultaneously recorded brain activity, we show that viewing behavior explains activity in a large network of regions and that DeepMReye can replace camera-based eye tracking for studying or accounting for these effects. The approach and results presented below emphasize the importance of eye tracking for MRI research and introduce a software solution that makes camera-less MR-based eye tracking widely available for free.

### Decoding gaze position from the eyeballs using convolutional neural networks

We demonstrate the wide applicability of our CNN-approach (Fig. 1AB) by decoding gaze from multiple existing fMRI datasets with a total of 268 participants performing diverse viewing tasks (Fig. 1D) including fixation (dataset 1, Alexander et al., 2017), smooth pursuit (dataset 2-4, Nau et al., 2018a, 2018b), visual search (dataset 5, Julian et al., 2018) and free picture viewing(part of dataset 6). These datasets were acquired on five 3T-MRI scanners using 14 scanning protocols. Repetition times (TR) ranged between 800-2500ms and voxel sizes ranged between 1.5-2.5mm. The eyeballs of each participant were first co-registered non-linearly to those of our group-average template, which was obtained by averaging the functional images of all participants in dataset 4 (Nau et al., 2018a) fixating at the screen center. For each participant, we first aligned the head, then a facial bounding box and finally the eyeballs to the ones of our template. This three-step procedure ensured that the eyeballs were aligned across participants and that the average gaze position reflected center fixation. The template brain has itself been co-registered to an MNI-structural template in which the eyes were manually segmented (Fig. 1A). We then extracted the multi-voxel-pattern (MVP) of the eyes at each imaging acquisition, normalized the pattern in time and space (Fig. 1B) and fed it into the CNN (Fig. 1C). While the exact model training and test procedure will be explained in detail later, it essentially uses the MVP of the eyes to predict 10 on-screen gaze coordinates corresponding to the respective volume. For the main analyses, these 10 gaze labels per TR were obtained either using camera-based eye tracking in case of the unconstrained visual search dataset (Julian et al., 2018), or from the screen coordinates of the fixation target in case of all others (Alexander et al., 2017; Nau et al., 2018a, 2018b). For the final model evaluation, these 10 gaze labels were median-averaged to obtain one gaze position per TR. The CNN was trained using cross-validation and a combination of two weighted loss functions (Fig. 1C): 1) the ‘Euclidean error’ between real and predicted gaze position and 2) a ‘predicted error’. The latter represents an unsupervised measure of the expected Euclidean error given the current input data.

**Figure 1:**
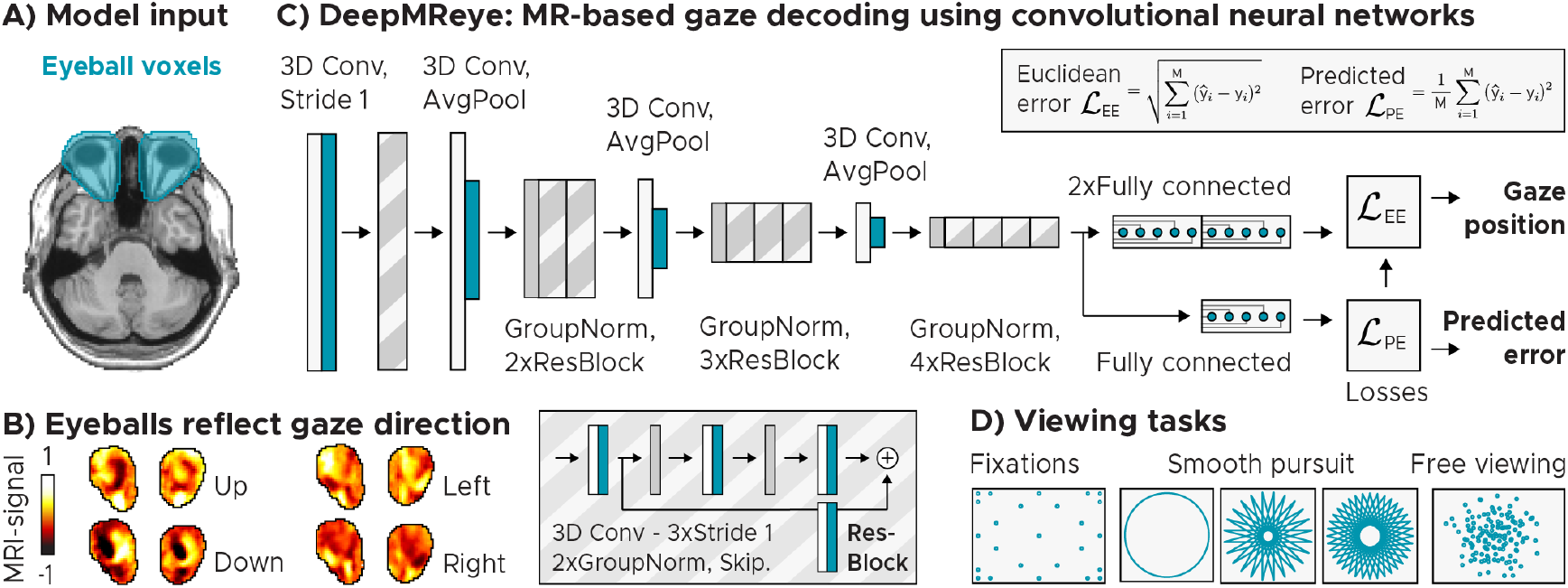
Model architecture and input. A) Manually delineated eye masks superimposed on T1-weighted structural template (Colin27)at MNI-coordinateZ = −36. B) Eyeball MR-signal reflects gaze direction. We plot the normalized MR-signal of eye-mask voxels of a sample participant who fixated a target on the left (X,Y = −10,0°), right (10,0°), top (0, 5.5°) or bottom (0, −5.5°) of the screen. C) Convolutional neural network architecture. The model takes the eye-mask voxels as 3D-input and predicts gaze position as a 2D (X, Y) regression target. It performs a series of 3D-convolutions (3D Conv) with group normalizations (GroupNorm)and spatial downsampling via average pooling (AvgPool) in between. Residual blocks (ResBlock) comprise an additional skip connection. The model is trained across participants using a combination of two loss functions: 1) The Euclidean error between the predicted and the true gaze position, and 2) the error between the Euclidean error and a predicted error. It outputs gaze position and the predicted error as a decoding-confidence measure for each TR. D) Schematics of viewing priors. We trained and tested the model on data of 268 participants performing fixations (Alexander et al., 2017), smooth pursuit on circular or star-shaped trajectories (Nau et al., 2018a, 2018b) and free viewing (Julian et al., 2018).

### Decoding viewing behavior in held-out participants

First, we examined the decoding performance in five key datasets that were acquired for other purposes (datasets 1-5, see Methods, Fig. 2, Alexander et al., 2017; Nau et al., 2018a, 2018b;Julian et al., 2018). The model was trained and tested using an across-participant decoding scheme, meaning that it was trained on 80% of the participants within each dataset and then tested on the held-out 20% of participants of that dataset. This procedure was cross-validated until all participants were tested once. For all viewing behaviors we found that the decoded gaze path followed the ground truth gaze path closely in the majority of participants (Fig. 2A). To quantify gaze decoding on the group level, we computed three measures: the Euclidean error (EE, Fig. Fig. 2B, S1), the Pearson correlation (r, Fig. 2C) as well as the coefficient-of-determination (*R*^2^, Fig. S2A) between the real and the decoded gaze paths of each participant. We found that gaze decoding worked in the large majority of participants with high precision (Fig. 2C, Fig. S2B) and for all viewing behaviors tested (Median performance of the 80% most reliable participants (low predicted error): All datasets: [r = 0.89, *R*^2^ = 0.78, EE = 1.14°], Fixation: [r = 0.86, *R*^2^ = 0.74, EE = 2.89°], Pursuit 1: [r = 0.94, *R*^2^ = 0.89, EE = 0.64°], Pursuit 2: [r = 0.94, *R*^2^ = 0.88, EE = 1.14°], Pursuit 3: [r = 0.86, *R*^2^ = 0.72, EE = 1.11 °], Free viewing: [r = 0.89, *R*^2^ = 0.78, EE = 2.17°]). These results were robust also when independent data partitions of each participant were used for training and test (within-participant decoding scheme, Fig. S4A), and that DeepMReye uncovered gaze position even when independent datasets were used for model training and test (across-dataset decoding, Fig. S4B). Together, these results demonstrate that gaze decoding with DeepMReye can be highly reliable and accurate. It allows reconstructing even complex viewing behaviors in held-out participants and detects outliers in an unsupervised fashion. Critically, it does so by relying solely on the MR-signal of the eyeballs without requiring any MR-compatible camera equipment.

**Figure 2:**
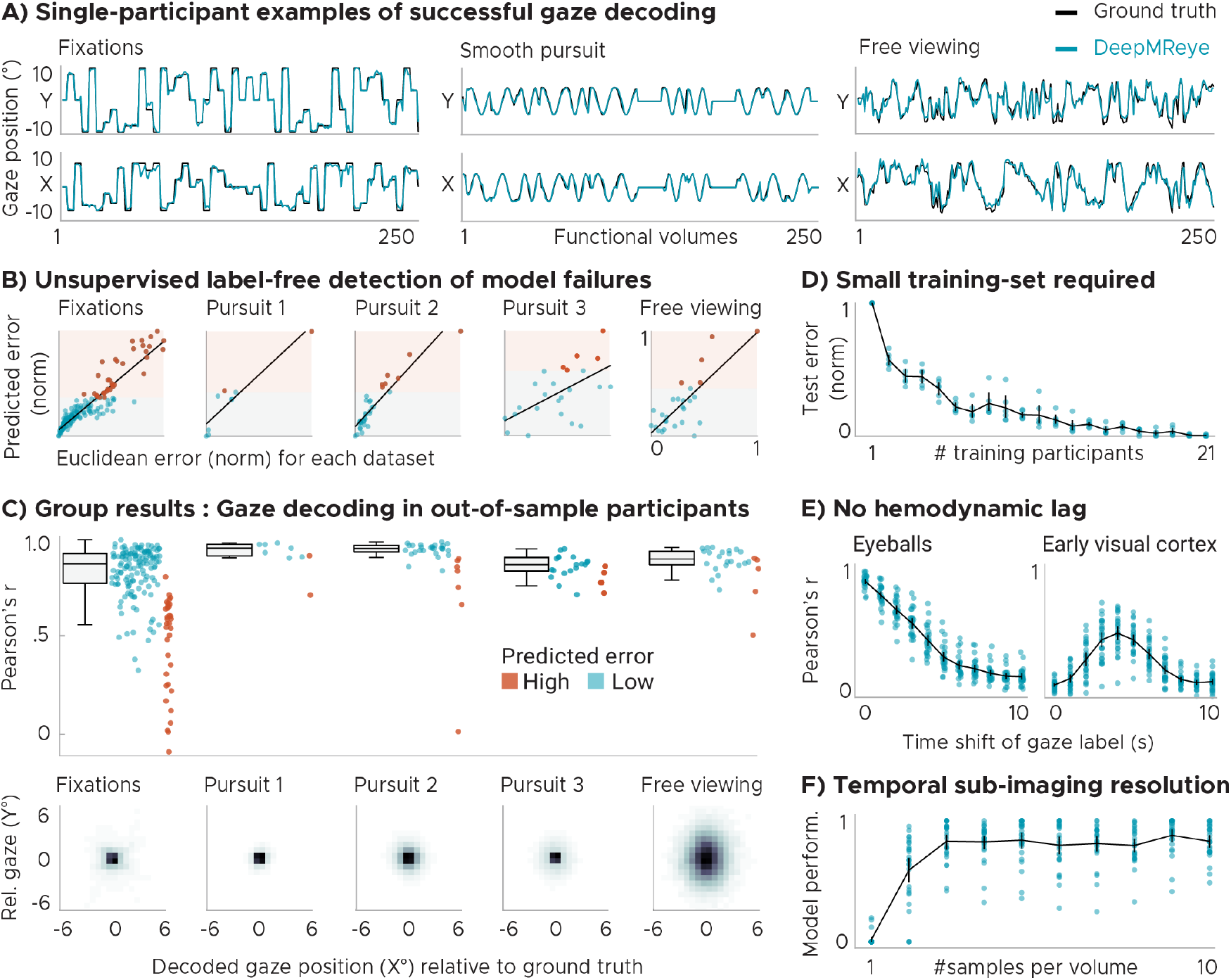
Across-participant gaze decoding results. A) Single-participant examples of successful gaze decoding for three viewing behaviors. B) Predicted error (PE) correlates with the Euclidean error between real and predicted gaze positions. This allows filtering the test set post-decoding based on estimated reliability. We plot single-participant data with regression line. Participants were split into 80% most reliable (Low PE, blue) and 20% least reliable participants (high PE, orange). Scores normalized for visualization. C) Group results: Top panel shows gaze decoding expressed as the Pearson correlation between true and decoded gaze trajectory for the five key datasets featuring fixations, 3x smooth pursuit and visual search. Participants are color coded according to PE. We plot Whisker-box-plots for Low-PE participants and single-participant data for all. Bottom panel shows time-collapsed group-average histograms of decoded positions relative to the true positions [0,0] in visual degrees. Color depicts decoding probability (black = high). D) Test error as a function of how many participants were used for model training. E) Gaze decoding from the eyeballs and early visual cortex for time-shifted gaze labels. F) Sub-imaging temporal resolution: We plot the model performance (explained variance normalized for each participant) depending on how many sub-imaging samples were decoded. D-F show results for visual search dataset 5.

### Unsupervised outlier detection

As mentioned above, the model computes a predicted error score for each sample and participant in addition to decoding gaze position. Importantly, this predicted error correlated with the true Euclidean error across participants, allowing to detect participants for which the decoding did not work well (Fig. 2B, Fig. S1AB). It can thus be used to remove outliers from subsequent analysis or to account for them for example by adding covariate regressors in group analyses. Note that besides detecting outlier participants, the predicted error also allowed removing outlier-samples within each participant, which further improved the reliability of the results (Fig. S3).

### No camera required for model training

We next explored our model’s requirements and boundary conditions in detail. First, we tested what type of training labels are required for DeepMReye, finding that both the screen coordinates of a fixation target (Fig. 2C) and labels obtained using camera-based eye tracking (Fig. S5) led to similar performance. While the results presented for dataset 5 (Fig. 2C) already reflect the ones obtained with camera-based labels, we additionally re-ran the model on gaze labels obtained via camera-based eye tracking also for the smooth pursuit datasets 3-4 (Fig. S5). Thus, because DeepMReye can be trained on fixation-target labels only, and because it generalizes across participants (Fig. 2), users could acquire fMRI data for a few participants performing various fixation tasks, record the screen coordinates of the fixation target as training labels, train the model on these labels and then decode from all other participants. Upon publication, we will provide the code for an experimental paradigm that can be used to produce such training labels (see ‘Data and code availability’ statement and ‘User recommendation’ section).

### Small training set

Next, we asked how many participants were required for model training. We tested this by iteratively sub-sampling the number of participants in the training set, each time testing how well the model performed on the same test participants. We chose to conduct this analysis on the data of dataset 5 because it featured the most natural and hence most complex viewing pattern tested. We found that model performance improved with an increasing training set size, but also that model performance already reached a ceiling level at as few as 6-8 participants (Mean performance, 1 participant: [r = 0.43, *R*^2^ = 0.11, EE = 5.12°], 5 participants: [r = 0.81, *R*^2^ = 0.62, EE = 3.18°], 10 participants: [r = 0.86, *R*^2^ = 0.71, EE = 2.58°], Fig. 2D, Fig. S6). This suggests that even a small training set can yield a well-trained model and hence reliable decoding results. Model performance likely also depends on how much data is available for each participant and on how similar the expected viewing behavior is between training and test set. If the gaze pattern is very similar across participants, which can be the case even for viewing of complex stimuli such as real-world scenes (Ehinger et al., 2009), decoding it in independent participants can work even better despite a small training set. This fact can be seen for example in our main results for the smooth-pursuit dataset 2 (Nau et al., 2018b, Fig. 2).

### No hemodynamic component

Naturally, when the eyes move, the surrounding tissue undergoes dramatic structural changes, which are expected to affect the MR-signal acquired at that time. To test whether this is the source of information used for decoding, we shifted the gaze labels relative to the imaging data by various TR’s (0-10), each time training and testing the model anew. Indeed, we found that the eyeball decoding was most accurate for the instantaneous gaze position and that no hemodynamic factors needed to be considered (Fig. 2E). This is in stark contrast to decoding from brain activity for which the same model pipeline can be used (Fig. 2E). In V1, decoding was optimal after around 5-6 seconds (r=0.483 ± 0.132) and followed the shape of the hemodynamic response function (HRF).

### Sub-imaging temporal resolution

Intriguingly, because different imaging slices were acquired at different times and because the MR-signal of a voxel can be affected by motion, it should in principle be possible to decode gaze position at a temporal resolution higher than the one of the imaging protocol (sub-TR resolution). As mentioned above, DeepMReye classifies 10 gaze labels per functional volume, which are median-averaged to obtain one gaze position per TR. This procedure yielded a higher decoding performance compared to classifying only one position, and it enabled testing how well the gaze path can be explained by the sub-TR labels themselves (Fig. S8A). We found that during visual search more gaze-path variance was explained by decoding up to three positions per TR compared to decoding only one position per TR (3Hz, Fig. 2F), which dovetails with the average visual-search eye-movement frequency of 3Hz (Wolfe, 2020). Moreover, the 10 real and decoded sub-TR labels varied similarly within each TR (Fig. S8B), which again suggests that within-TR movements could be detected. While the exact resolution likely depends on the viewing behavior and the imaging protocol, these results show that at least a moderate sub-imaging temporal decoding resolution is indeed feasible.

### Across-dataset generalization

The results presented so far show that the gaze decoding with DeepMReye is highly accurate when the viewing behavior and the imaging protocol are similar between training and test set. To test if our model also generalizes across datasets, we next implemented a leave-one-dataset-out cross-validation scheme. Most datasets were acquired by different groups using different MR-scanners, participants and viewing behaviors but with similar voxel sizes and TR’s. While this across-dataset scheme led to overall lower performance scores compared to the across-participant (within-dataset) scheme presented earlier, it nevertheless recovered viewing behavior with remarkable accuracy in all cases (Median performance of the 80% most reliable participants (low predicted error): All datasets: [r = 0.84, *R*^2^ = 0.59, EE = 2.78°], Fixation: [r = 0.79, *R*^2^ = 0.52, EE = 5.34°], Pursuit 1: [r = 0.88, *R*^2^ = 0.64, EE = 1.47°], Pursuit 2: [r = 0.86, *R*^2^ = 0.65, EE = 2.15°], Pursuit 3: [r = 0.85, *R*^2^ = 0.55, EE = 2.01 °], Free viewing: [r = 0.84, *R*^2^ = 0.61, EE = 2.96°], Fig. S4). This suggests that datasets acquired with similar fMRI protocols can be used for model training, even if the recording site or the protocol were not exactly the same. Future investigations will need to quantify how larger differences in scan parameters affect this across-dataset generalization (e.g. different phase encoding directions or slice tilts). Note that despite higher Euclidean error and lower *R*^2^-scores compared to within-dataset decoding, the across-dataset decoding scheme led to relatively high Pearson correlations. This indicates that the main reason for the lower performance scores is the scaling of the decoding output relative to the test labels, likely because the data range of the training and testing labels differed. Importantly, this also suggests that the presence of putative eye movements, but not their correct amplitude, could still be detected accurately, which is the most important aspect for many fMRI analyses or nuisance models.

### Robust across voxel sizes and repetition times

Functional MRI protocols can differ in many aspects. Most importantly in this context, they can differ in the spatial and temporal resolution of the acquired data (i.e. voxel size and TR). To explore the influence of these two parameters on the decoding performance in detail, we varied them systematically across 9 fMRI protocols for the acquisition of a sixth dataset. For each of the 9 sequences, we scanned 4 participants with concurrent camera-based eye tracking while they freely explored pictures (Hebart et al., 2019) or performed fixation (Alexander et al., 2017) and smooth pursuit tasks similar to the ones used earlier (Nau et al., 2018a, 2018b). DeepMReye decoded gaze position robustly in this dataset 6 during all of these tasks and in all imaging protocols tested (3×3 design: TR = 1.25s, 1.8s, 2.5s, voxel size = 1.5mm, 2mm, 2.5mm, Fig. 3A), demonstrating that it is widely applicable across a broad range of routinely used voxel sizes and TR’s.

**Figure 3:**
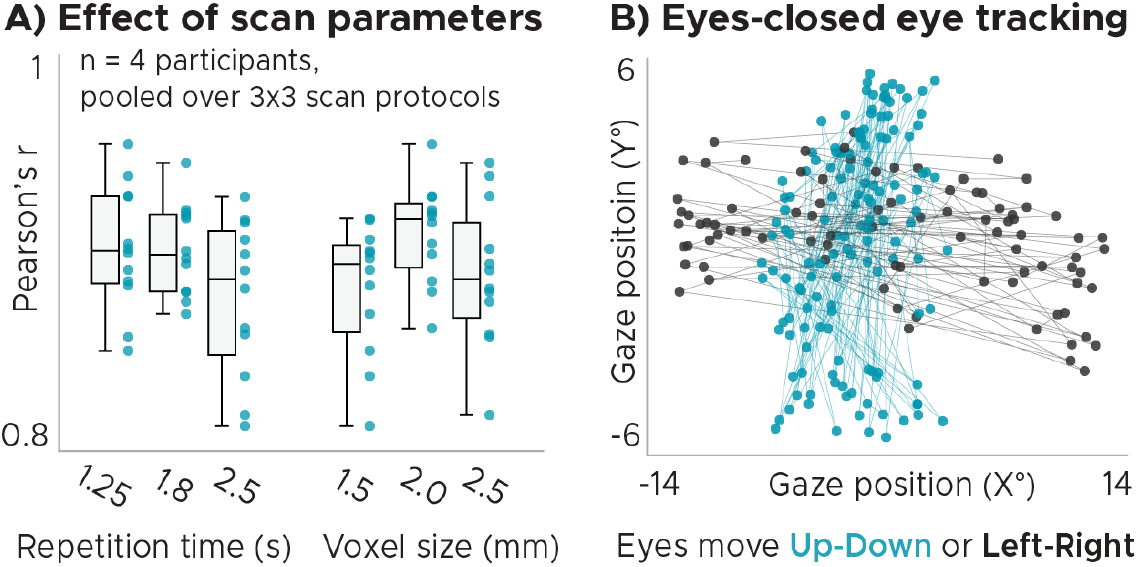
Effect of scan parameters and eye tracking while the eyes are closed. A) Effect of voxel size and repetition time (TR). We plot gaze decoding expressed as the Pearson correlation between true and decoded gaze trajectory for different voxel sizes and TR’s. We plot Whiskerbox-plots and single-participant data (n = 4) for 9 fMRI protocols collapsed either over TR or voxel size. DeepMReye recovered viewing behavior successfully in all sequences tested. B) Decoded gaze coordinates for a participant being instructed to move the eyes left & right or up & down while keeping them closed. Dots are colored based on button press of participant indicating movement direction.

### Eyes-closed tracking

Traditional MR-compatible eye-tracking systems typically detect certain features of the eyes such as the pupil and/or the corneal reflection in a video, which are then tracked over time (Duchowski, 2017). When the relevant features are occluded or cut off on the video (e.g. when the eyes close), the tracking is lost. Because our approach relies on the fact that the eyeball MR-signal changes as a function of gaze position (Fig. 1B), it might be possible to decode gaze position, or in this case more generally the state of the eyeballs, even when the eyes are closed. As a proof-of-concept, we therefore tested in one participant of dataset 6 whether DeepMReye can uncover viewing behavior even when the eyes are closed. The participant was instructed to close the eyes and move them either repeatedly from left to right or from top to bottom, and to indicate the behavior via key press. We trained DeepMReye on the diverse eyes-open viewing data from all participants in dataset 6 and then decoded from the one participant while the eyes were closed. We found that the gaze pattern decoded with DeepMReye closely matched the participant’s self-report, suggesting that it is indeed possible to perform eye tracking while the eyes are closed (see the ‘User recommendation’ section).

### Viewing behavior explains network-wide brain activity

The results presented so far demonstrate that DeepMReye can be used to perform eye tracking in many experimental settings. A critical open question that remained was whether its decoding output can be used to analyze brain activity. To test this, we implemented a whole-brain mass-univariate general model (GLM) for the visual search dataset 5. We again chose this dataset because it featured the most complex viewing pattern tested. To simulate differences in viewing behavior between the two conditions, we first computed an eye-movement index, reflecting the Euclidean distance between gaze positions of subsequent volumes. We used this eye-movement index to build two main regressors of interest, one modeling large eye movements and one modeling short eye movements. Both regressors were binarized and convolved with the hemodynamic response function. Contrasting the model weights estimated for these two regressors was expected to reveal regions in the brain whose activity is driven by viewing behavior such as the visual and oculomotor (attention) network (Berman et al., 1999; Petit & Haxby, 1999).

To know what we were looking for, we first conducted this analysis using the gaze labels obtained with traditional camera-based eye tracking and then compared the results to the ones obtained for the three cross-validation schemes of DeepMReye (within-participants, across-participants, across-datasets).

As predicted, we found that viewing behavior explained brain activity in a large network of regions (Fig. 4) including the early visual cortex, frontoparietal regions (likely the frontal eye fields), the posterior parietal cortex as well as temporal lobe regions (likely including the human motion complex). Importantly however, differences in viewing behavior also explained brain activity in regions not typically associated with oculomotor function such as the ventromedial prefrontal cortex (vmPFC), the anterior and posterior cingulate cortex, the medial parietal lobe (likely comprising the retrosplenial cortex), the parahippocampal gyrus as well as the hippocampus (Fig. 4).

**Figure 4:**
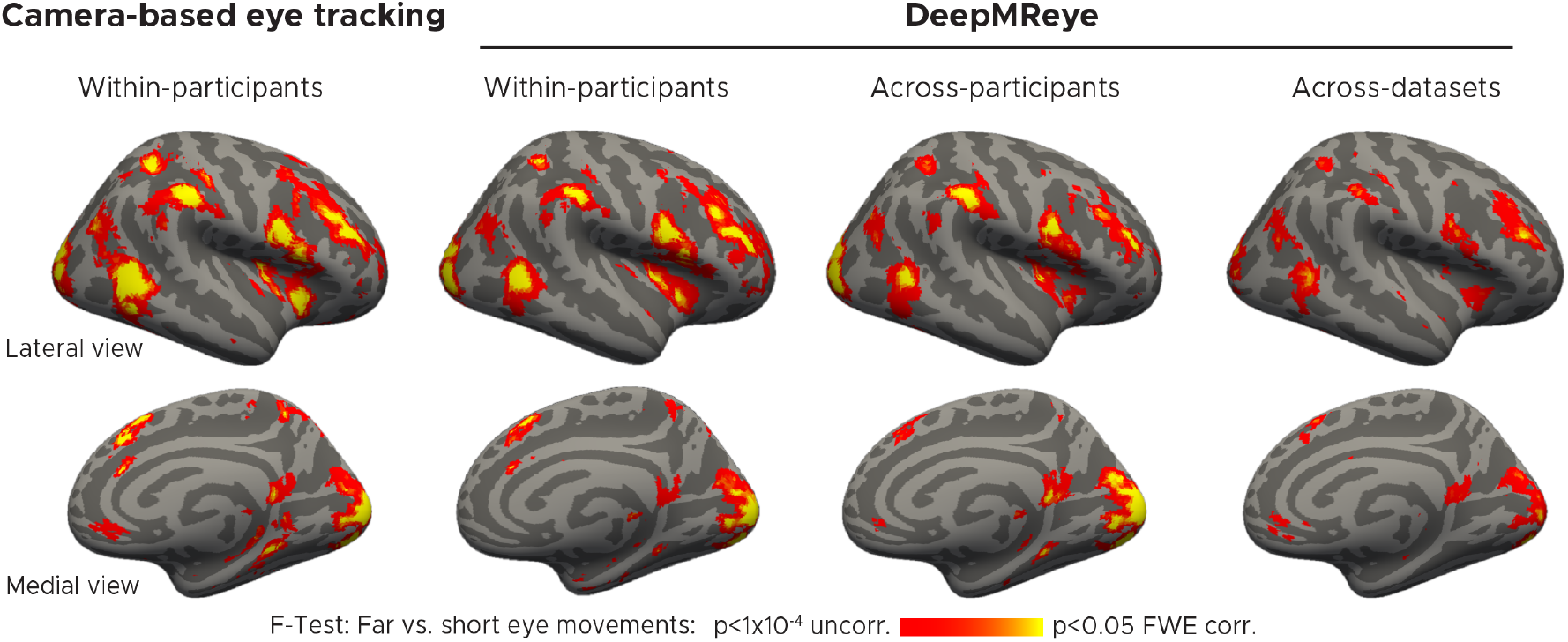
Decoded viewing behavior explains network-wide brain activity. General-linear-model (GLM) group results for the contrast ‘Far vs. short eye movements’ during visual search. We plot the F-statistic of this contrast superimposed on a templatesurface (fsaverage) for gaze-labels obtained with camera-based eye tracking (first panel) as well as for three DeepMReye cross-validation schemes. Within-participants: All participants of a dataset were included with different partitions in model training and test. Across-participants: Different participants were included during model training and test. Across-datasets: Different datasets (and hence also different participants) were included during model training and test.

Strikingly, comparing the results obtained with DeepMReye to the ones obtained with camera-based eye tracking showed an exceptional match between the two (Fig. 4). This was true for all decoding schemes, including the across-participant decoding scheme, which can be conducted even in existing datasets with some preparation (Fig. 2, see ‘User recommendations’). Moreover, even the across-dataset scheme explained gaze related variance on group level, despite the differences in the underlying viewing behaviors and imaging protocols.

Finally, because eye movements are associated not only with brain activity but also with imaging artifacts, the MRI signal might also be affected instantaneously when the movement occurs. To quantify these instantaneous effects, we repeated the GLM analysis modeling eye-movement related fluctuations in the MRI signal without accounting for the hemodynamic response. This variance is not captured by traditional head-motion regressors (Fig. S9). Again, we found that eye movements explained signal variations in many brain regions (Fig. S10), likely reflecting a combination of imaging artifacts and instantaneous hemodynamic components (e.g. the initial dip).

## Discussion

DeepMReye is a camera-less eye tracking framework based on a CNN that decodes gaze position from the MR-signal of the eyeballs. It allows monitoring viewing behavior accurately and continuously at a moderate sub-imaging resolution without the need for MR-compatible cameras. We demonstrated that our approach works robustly for a wide range of voxel sizes and repetition times as well as for various viewing behaviors including fixation, smooth pursuit, free viewing and as a proof-of-concept even when the eyes were closed. For each gaze position and participant, the model outputs an unsupervised predicted error score that can be used to filter out outliers even when test labels are missing. A small training set can yield a well-trained model and high decoding performance even when trained without camera-based labels. The decoded gaze positions and eye movements can be used in subsequent fMRI analyses similar to camera-based eye tracking, and doing so here revealed gaze-related activity in a large network of regions in the brain (Berman et al., 1999; Petit & Haxby, 1999; Voss et al., 2017). Critically, by testing our model in independent participants within each dataset, but also in participants in other datasets acquired with different MR-scanners and protocols, we demonstrated the potential of DeepMReye to successfully decode viewing behavior also in existing fMRI data.

### MR-based gaze prediction

The present work is directly inspired by earlier reports showing that the MR-signal of the eyeballs can be used to infer the state of the eyes during MRI-scanning. This includes movements of the eyes (Tregellas et al., 2002; Beauchamp, 2003; Keck et al., 2009; Franceschiello et al., 2020), the position of gaze on the screen (Heberlein et al., 2006; LaConte & Glielmi, 2006; Son et al., 2020; Sathian et al., 2011; Keck et al., 2009) or whether the eyes were open or closed (Brodoehl et al., 2016). Moreover, gaze position can be decoded from early visual cortex activity during scene viewing (O’Connell & Chun, 2018) and as shown here during visual search (Fig. 2E). However, DeepMReye goes beyond these earlier reports in multiple ways. Most importantly, earlier approaches such as predictive-eye-estimation-regression (PEER, Son et al., 2020) required calibration data for every single participant, meaning that at least two calibration scans need to be acquired during each scanning session. In contrast, our deep-learning based approach generalizes across participants, allowing to perform eye tracking even when training and test labels are missing. The model could be trained on the data of a few participants and then used for decoding from all other participants. Moreover, earlier approaches were limited to the sampling resolution of the imaging protocol, meaning that one average gaze position per functional image could be extracted. In contrast, we extracted gaze position at a moderate sub-TR resolution (~3Hz) and with higher accuracy than previous approaches, allowing to perform MR-based eye tracking with a higher level of detail. Third, as a proof-of-principle, we show that our model reconstructs viewing behavior even when the eyes are closed. Finally, we provide the first open source and user-friendly implementation for MR-based eye tracking as an interactive decoding pipeline inspired by other fMRI open source initiatives (e.g. (Esteban et al., 2019)). DeepMReye hence overcomes several critical limitations of earlier work, presenting the most general and versatile solution to camera-less eye tracking in MRI to date.

### What information does the model use?

Eye movements naturally entail movements of the eyeballs but also of the optic nerves and the fatty tissue around them. To capture these movements, our custom eye masks cover a large area behind the eyes excluding skull and brain tissue. When the eyes move, the multi-voxel-pattern in these masks changes drastically (Fig. 1B), an effect that might be even amplified by the magnetic field distortions often occurring around the eyes. DeepMReye hence likely utilizes information traditionally considered to be motion artifacts, which are not corrected by classical realignment during preprocessing (Fig. S9, Fig. S10). The fact that the actual motion of the eye is used for decoding also means that no hemodynamic lag needs to be considered (Fig. 2E). The current gaze position is decoded directly from each TR respectively. We believe that two sources of information further contribute to the moderate sub-imaging decoding resolution that we observed. First, different imaging slices are being acquired at a different time within each TR and thus inherently carry some sub-TR information. This is true also for fMRI protocols that use multiband acquisition, which includes all datasets tested here. Future studies could examine the effect of slice timing on the decoding resolution in more detail. Second, similar to motion blur in a long-exposure camera picture, the MR-signal intensity of a voxel can itself be affected by movements. The multi-voxel-pattern at each TR might hence reflect how much the eyes moved, and the same average gaze position might look different depending on which positions were sampled overall within the respective TR.

### Looking forward

DeepMReye offers a multitude of exciting applications ranging from simple behavioral monitoring over confound removal to new and improved task-based analyses. Most basically, it offers an additional and low-effort behavioral read-out for any fMRI-experiment and allows to monitor task compliance for example by verifying that a fixation cross was fixated. Removing samples at which fixation was not maintained from subsequent analysis has been shown to improve predictive modeling results (LaConte & Glielmi, 2006) and may help to reduce the effects of in-scanner sleep more easily (Tagliazucchi & Laufs, 2014).

Our approach enables studies of the relationship between viewing and brain activity, and may more generally be used to inform almost any type of task-based model about the underlying viewing behavior. This could for example further improve the explanatory power of predictive models (Naselaris et al., 2011; Kriegeskorte & Douglas, 2019), and be especially promising for naturalistic free-viewing paradigms because the currently attended aspect of a stimulus can be taken into account (Sonkusare et al., 2019).

Importantly, eye movements can also be a major source of confounds in neuroimaging studies. As mentioned in the introduction, if differences in viewing between two conditions remain undetected, the interpretation of neuroimaging results may be compromised. We demonstrated here that many brain regions are affected by this issue, many of which are not typically studied in the context of eye movements (Fig. 4). Moreover, eye movements are associated with imaging artifacts that can affect data integrity throughout the brain (McNabb et al., 2020). A popular way of minimizing such confounds is having participants fixate at a fixation cross, which is helpful but also puts artificial constraints on a behavior that is fundamental to how we explore the world. Moreover, task compliance cannot always be guaranteed. DeepMReye may allow to identify and potentially compensate such confounds and artifacts for example by adding eye movement regressors directly to a GLM analysis as it is standard practice for head-motion regressors. This promises to improve the interpretability of task-based and resting-state fMRI results alike because nuisance variance would no longer be assigned to the regressors of interest (Murphy et al., 2013).

Thus, DeepMReye can provide many experimental and analytical benefits that traditional eye-tracking systems can provide too. Critically, it does so without any expensive equipment, trained staff or experimental time to be used. It can therefore be used widely in both research and clinical settings for example to study or diagnose neurodegenerative disorders (Anderson & MacAskill, 2013). Excitingly, it can even go beyond traditional eye tracking in certain aspects, offering new experimental possibilities that cannot easily be realized with a camera. For example, eye movements can be tracked even while the eyes are closed, suggesting it could be used to study oculomotor systems in the total absence of visual confounds, during resting state, and potentially even during rapid eye movements (REM) sleep. Moreover, the across-participant generalization enables new studies of patient groups for which camera-based eye trackers are not applicable. For example, DeepMReye could be trained on data of healthy volunteers and then tested on visually impaired participants for whom camera-based eye trackers cannot be calibrated. Most importantly, it allows gaze decoding in already existing task-based and resting-state fMRI datasets, in principle including all datasets that comprise the eyeballs. It could hence make new use of a large, existing and instantly available data resource (see “User recommendations”).

Finally, the same model architecture can be used to decode gaze position not only from the eyeballs but also from brain activity directly. Doing so is as simple as replacing the eye masks by a regions-of-interest maskof a certain brain region and accounting for the hemodynamic lag. We demonstrated this possibility using fMRI data from area V1 (Fig. 2E). Likewise, the same decoding pipeline could be used to decode other behavioral or stimulus features from brain activity, again showing the power of deep-learning-based methods for image analysis and neuroscience in general (Frey et al., 2019; Shen et al., 2017).

### Limitations

It is important to note that DeepMReye also has certain limitations and disadvantages compared to camera-based eye tracking. First, the eyeballs need to be included in the MRI images. This may not always be possible and can affect the artifacts that eye movements can induce. In practice, however, many existing and future datasets do include the eyes, and even if not, DeepMReye could still be used to decode from brain activity directly. Second, despite decoding at a temporal resolution that is higher than the one of the underlying imaging protocol, our approach does by no means reach the temporal resolution of a camera. Many aspects of viewing behavior happen on a time scale that can hence not be studied with DeepMReye. For experiments requiring such high temporal resolution, for example for studying individual saccades, we therefore recommend a camera system. However, many fMRI studies will not require monitoring gaze at high temporal resolution. This is because the regression analyses that are most commonly used in neuroimaging require the eye-tracking data to be downsampled to the imaging resolution irrespective of the sampling rate at which they were recorded. This means that even if gaze behaviorwas monitored at 1000 Hzwith a camera, the effective eye-tracking data resolution that enters the fMRI analysis is often the same as the one of DeepMReye. Also, many MRI facilities simply do not have an MR-compatible camera, leaving MR-based eye tracking as the only available option.

## Conclusions

In sum, DeepMReye is a camera-less deep-learning based eye tracking framework for fMRI experiments. It works robustly across a broad range of gaze behaviors and imaging protocols, allowing to reconstruct viewing behavior with high precision even in existing datasets. This work emphasizes the importance and the potential of combining eye tracking and neuroimaging for studying human brain function and provides a user-friendly and open source software solution that is widely applicable post-hoc.

## Author Contributions

MF & MN conceptualized the present work, developed the decoding pipeline and analyzed the data with input from CFD. MF wrote the key model implementation code with help from MN. MN acquired most and analyzed all datasets, visualized the results and wrote the manuscript with help from MF. MF, MN and CFD discussed the results & contributed to the manuscript.

## Data and code availability

Upon publication, we will share online our model code, documentation and Colab notebooks as well as eye-tracking calibration scripts that can be used to acquire training data for DeepMReye. In addition, we share our pre-trained model weights estimated on all datasets used in the present work. These model weights allow decoding viewing behavior without re-training the model in certain scenarios (see “User recommendation” section for details). All shared code will be available here: https://github.com/CYHSM/DeepMReye.

## Acknowledgements

We thank Ignacio Polti, Joshua B. Julian, Russell Epstein and Andreas Bartels for providing imaging and eye-tracking data that was used in the present work. We further thank Caswell Barry for helpful discussions and Joshua B. Julian and Christopher I. Baker for comments on an earlier version of this manuscript. This work is supported by the European Research Council (ERC-CoG GEOCOG 724836). CFD’s research is further supported by the Max Planck Society, the Kavli Foundation, the Centre of Excellence scheme of the Research Council of Norway – Centre for Neural Computation (223262/F50), The Egil and Pauline Braathen and Fred Kavli Centre for Cortical Microcircuits and the National Infrastructure scheme of the Research Council of Norway – NORBRAIN (197467/F50).

## User recommendations

### General recommendations

Despite successfullyapplying DeepMReye to data obtained with 14 scanning protocols, our datasets still capture only a limited number of sequence parameters and behaviors. We therefore generally recommend running a pilot study using the setup and the imaging protocol that was or will be used in the acquisition of the to-be-analyzed dataset. The larger the sample size used for model training, the more data is being acquired for each participant, and the more similar the viewing behavior is to the one of the test set, the better the decoding will be.

We further recommend thinking about whether the dataset includes at least some moments at which ground truth positions are known. For future studies, such scenarios could be added by design to validate the decoding output later on e.g. by presenting a fixation cross at various locations on the screen in the course of scanning. If the viewing behavior in the test set is unknown, we recommend training the model on a mixture of smooth pursuit and fixation with variable duration while sampling as many screen locations as possible.

To ensure that the eye masks fit every participant, the input data is being warped into our own functional MNI group template space. Many popular normalization algorithms rely on brain tissue segmentation for normalization and hence typically neglect the eyeballs. We therefore recommend users to apply non-linear warping algorithms that are agnostic to the underlying tissue. Here, we used the Advanced Normalization Tools (ANTs) running under Python.

Note that the placement of the mirror inside the MRI has a large impact on the position of the eyeballs relative to the screen. Even if participants fixate at the same position on the screen, depending on the mirror placement the eyeballs might be oriented differently. Our three-step co-registration procedure of the eyes mitigates this problem, but we still recommend users who acquire new data to place the mirror at approximately the same location and angle for all participants.

For users wishing to perform eye tracking while the eyes are closed, we recommend training the model on a combination of eyes-closed and eyes-open gaze labels. One option to obtain such labels would be to have participants fixate at various target locations on the screen and then close their eyes without moving them for 2-3 s. For studies planning on using DeepMReye during in-scanner sleep, we further recommend adding an awake but eyes-closed validation paradigm to the study, which could be similar to the one used here (Fig. 3B).

Finally, irrespective of which of the following decoding option is used, we recommend assessing the predicted error scores carefully for each participant (Fig. 2B, Fig. S1). The predicted error score is tightly correlated with the Euclidean error between real and decoded gaze position and hence allows to detect and remove outliers even when test labels are missing.

In the following, we outline multiple ways of how DeepMReye can be used. Which option is best depends on how much experimental time and data are available.

### User option 1

To maximize accuracy and robustness, we recommend users who acquire new data to scan a short calibration paradigm in addition to their regular experimental paradigm. This could be as simple as presenting various fixation targets sequentially at different locations on the screen (see e.g. dataset 1). Upon publication, we will provide the code for such a calibration scan online (see’Data & code availability’ statement). Importantly, the more similar the viewing behavior is between training and test set the more accurate the decoding will be. Acquiring such calibration data will allow training the model on data acquired in the same participants with the same fMRI sequence that the model is being tested on, which promises the best result. This option is recommended in any case, but especially when eyes-closed data are being analyzed. Note that in this case the model is still evaluated on data of all participants, with the calibration data used for model training and the actual experimental data used for decoding.

### User option 2

For users who cannot scan individual eye tracking calibrations for all participant, we recommend acquiring such calibration data at least for a subset of participants for example during piloting. This will allow training the model on some participants scanned on the same MRI scanner with the same imaging protocol, and then decode from others. This also offers a solution for already existing datasets. In this case, we again recommend scanning calibration data for a few participants with the same imaging protocol on the same MRI scanner if possible. We suggest acquiring calibration data for at least 6-8 participants, and to carefully evaluate the model performance within the training set. If necessary, more participants can be added. In addition, stronger dropout regularization and/orthe use of ensemble learning with various dropout strengths can help to improve model training in case of small sample sizes.

### User option 3

If no new calibration data can be acquired, users may use an already pre-trained version of DeepMReye that we provide. Note that we did show that our model generalizes across datasets, but also that it performed less accurately than the across-participant prediction within each dataset. We therefore recommend this option only for datasets in which at least some ground-truth-validation labels are known. Having said that, if more training data are being added to the model training in the future, the across-dataset prediction is expected to further improve. Especially important will be data featuring even more diverse viewing behaviors and imaging protocols.

## Methods

### Datasets

DeepMReye was trained and tested on data of 268 participants acquired on five 3T MRI scanners with 14 different scanning protocols and various pre-processing settings. The individual datasets are described below and were partially used in earlier reports. For other details of each individual dataset please see the original published articles (Alexander et al., 2017; Nau et al., 2018a, 2018b; Julian et al., 2018). A T1-weighted structural scan with 1 mm isotropic voxel resolution was acquired for all participants, and camera-based eye-tracking data was included for participants in datasets 3-6. An overview of the datasets is provided in Table 1.

**Table 1:**
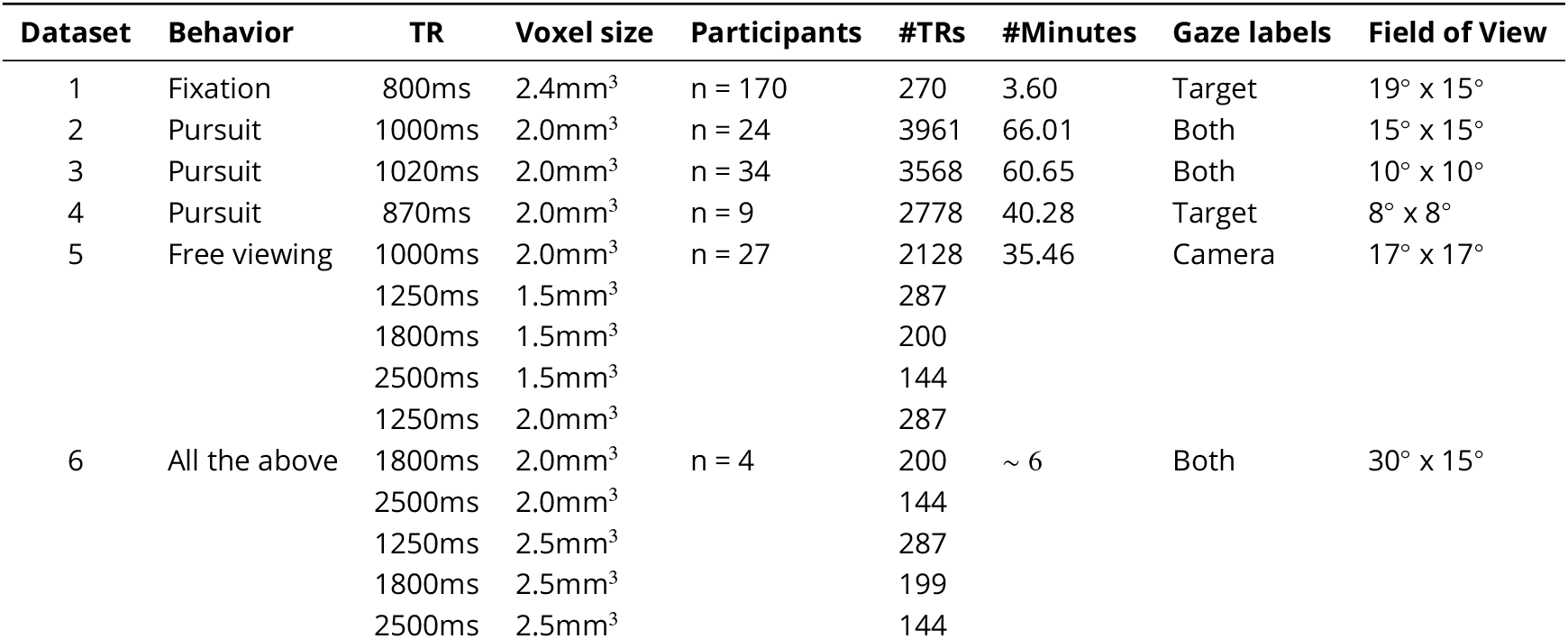
Overview of the six datasets. We list the dataset number, the viewing behavior that was tested, the repetition time (TR) and voxel size of the imaging protocol, the number of participants, the amount of data acquired for each participant expressed as the average number of acquired volumes (#TRs) and as the total scanning time (#Minutes), the type of gaze labels that DeepMReye was trained and tested on (incl. camera-based labels, screen coordinates of the fixation target, or both) as well as the task-relevant field of view (FoV) of the participant.

#### Dataset 1: Fixation and saccades

##### Data & task

These data were made publicly available by Alexander and colleagues (Alexander et al., 2017) and were downloaded from the Healthy Brain Network (HBN) Biobank (http://fcon_1000.projects.nitrc.org). These data were also used in earlier reports (Son et al., 2020). It is part of a larger and ongoing data collection effort with pediatric focus and comprises participants between 5 and 21 years of age. We limited our analysis to a subset of the full dataset for which we ensured that there were no visible motion artifacts in either the T1-or the average T2*-weighted images and the eyeballs were fully included in the functional images. We included 170 participants in total. For each participant, at least two fMRI runs were scanned in which they performed a typical eye tracking calibration protocol. In each run, they fixated at a fixation target that sequentially moved through 27 locations on the screen, with each location being sampled twice for 5s. Gaze positions were sampled within a window of X= 19° and Y = 15° visual angle. The screen coordinates of the fixation target served as training and testing labels for the main analyses (Fig. 2).

##### fMRI-data acquisition & preprocessing

Imaging data were acquired on a Siemens 3T Tim Trio MRI scanner located at the Rutgers University Brain Imaging Center, Newark, USA. Following EPI parameters were used: voxel size = 2.4mm isotropic, TR = 800ms, TE = 30ms, flip angle = 31 °, multiband factor = 6. Images were coregistered to our template space as described below.

#### Dataset 2: Smooth pursuit 1

##### Data & task

These data were used in one of our previous reports (Nau et al., 2018b). Nine participants performed a smooth pursuit visual tracking task in which they either tracked a fixation target moving on a circular trajectory with a radius of 8° visual angle or one that remained at the screen center. In addition, planar-dot-motion stimuli were displayed in the background moving on the same circular trajectory at various speeds. This resulted in a total of 9 different conditions. Following pursuit and motion speed combinations were tested in separate trials: [eye, background in °/s] = [0,0], [0,1], [0,3], [2,1], [2,2], [2,3], [3,2], [3,3], [3,4]. These conditions were tested in blocks of 12 seconds in the course of 34 trials over 4 scanning runs of ~10.5 minutes each. To balance attention across conditions, participants performed a letter-repetition-detection task displayed on the fixation target. Gaze positions were sampled within a window of X = 8° and Y = 8° visual angle. The screen coordinates of the fixation target served as training and testing labels for our model.

##### fMRI-data acquisition & preprocessing

Imaging data were acquired on a Siemens 3T MAGNETOM Prisma MRI scanner located at the Max-Planck-Institute for Biological Cybernetics in Tuebingen, Germany. Following EPI-parameters were used: voxel size = 2mm isotropic, TR = 870ms, TE = 30ms, flip angle = 56°, multiband factor = 4, GRAPPA factor = 2. Note that 9 other participants were excluded because the functional images did not or only partially included the eyeballs. Images were corrected for head motion and field distortions using SPM12 (www.fll.ion.ucl.ac.uk/spm/) and then coregistered to our template space as described below.

##### Eye tracking

We monitored gaze position at 60 Hz using a camera-based eye tracker by Arrington Research. Please note that these eye-tracking data showed a higher noise level than the other datasets due to drift and because the pupil was frequently lost. We therefore used the screen coordinates of the fixation target for model training and testing as in dataset 1. To still visually compare the decoding output to the eye-tracking data post-hoc, we removed blinks, detrended the eye tracking time series using a second-order polynomial function and median-centered it on the screen center. We removed samples in which the pupil was lost by limiting the time series to the central 14×14 degree visual angle, smoothed it using a running-average kernel of 100ms and scaled it to match the data range of the fixation target using the sum-of-squared-errors as loss function. The time series was then split into the individual scanning acquisitions.

#### Dataset 3: Smooth pursuit 2

##### Data & task

These data are currently being analyzed for another report and comprised 34 participants (Polti & Nau et al., in prep). Like in dataset 2, participants performed a smooth pursuit visual tracking task in which they fixated at a fixation target moving on a star-shaped trajectory. Twenty-four eye movement directions were sampled in steps of 15° at four speed levels: 4.2°/s, 5.8°/s, 7.5°/s and 9.1 °/s. Speeds were interleaved and sampled in a counterbalanced fashion. In addition to the visual tracking task, participants performed a time-to-collision (TTC) task. The trajectory was surrounded by a circular yellow line on gray background with a radius of 10° visual angle centered on the screen center. Whenever the fixation target stopped moving before switching direction, participants indicated by button press when the target would have touched the yellow line if it continued moving. Gaze positions were sampled within a window of X = 10° and Y = 10° visual angle. Each participant performed a total of 768 trials in the course of 4 scanning runs with 16-18 minutes (including a short break in the middle). The screen coordinates of the fixation target served as training and testing labels for the main analyses (Fig. 2).

##### fMRI-data acquisition & preprocessing

Imaging data were acquired on a Siemens 3T MAGNETOM Skyra located at the St. Olavs Hospital in Trondheim, Norway. Following EPI-parameters were used: voxel size = 2mm isotropic, TR = 1020ms, TE = 34.6ms, flip angle = 55°, multiband factor = 6. Images were corrected for head motion using SPM12. The FSL topup function was used to correct field distortions using an image acquired with the same protocol except that the phase-encoding direction was inverted (https://fsl.fmrib.ox.ac.uk/fsl/fslwiki/topup). Images were then coregistered to our template space as described below.

##### Eyetracking

We monitored gaze position during the experiment at a rate of 1000 Hz using an MR-compatible infrared-based eye tracker (Eyelink1000). Blinks were removed, the time series was downsampled to 100hz, linearly detrended within each scanning run and smoothed with a running-average kernel of 100ms. We then split the time series into individual scanning acquisitions (TR’s) to obtain the final training and testing gaze labels for our model. The camera-based eye tracking labels served as training and testing labels for supplementary analyses (Fig. S5).

#### Dataset 4: Smooth pursuit 3

##### Data & task

These data were used in one of our previous reports (Nau et al., 2018a). Twenty-four participants performed a smooth pursuit visual tracking task in which they tracked a fixation target moving at a speed of 7.5°/s on a star-shaped trajectory with 36 directions. The target moved within a virtual arena which participants oversaw from bird’s eye view. Eye movement directions were sampled in steps of 10°. In a visual-motion control condition, the target remained at the screen center and the arena moved instead. Participants additionally performed a spatial memory task. They memorized the location of colored objects on the screen, which were shown only when the fixation target moved across them. Gaze positions were sampled within a window of X = 15° and Y = 15° visual angle. Each participant performed a total of 81 trials in the course of 9 scanning runs. This included 54 smooth pursuit trials of 60 seconds each and 27 center fixation trials of 30 seconds each. The screen coordinates of the fixation target served as training and testing labels for the main analyses (Fig. 2).

##### fMRI-data acquisition & preprocessing

Imaging data were acquired on a Siemens 3T MAGNETOM PrismaFit MRI scanner located at the Donders Centre for Cognitive Neuroimaging, Nijmegen, the Netherlands. Following EPI-parameters were used: voxel size = 2mm isotropic, TR = 1000ms, TE = 34ms, flip angle = 60°, multiband factor = 6. Data were realigned using SPM12 (www.fll.ion.ucl.ac.uk/spm/) and coregistered to our template space as described below.

##### Eye tracking

Similar to dataset 3, we again monitored gaze position during the experiment at 1000 Hz using an Eyelink 1000 eye tracker. Blinks were removed and the data were downsampled to the monitor refresh rate of 60hz. We then reduced additional tracking noise by removing samples at which the pupil size diverged more than one standard deviation from the mean, by removing the inter-trial-interval during which most blinks occurred and by smoothing the time series with a running-average kernel of 100ms. We then linearly detrended and median-centered the time series of each trial individually to remove drift. Finally, we split the time series according to the underlying scanner acquisition times to create our final training and testing labels for this dataset. Note that the original dataset (Nau et al., 2018a) comprises 5 additional participants for which no eye-tracking data has been obtained and that were excluded. The camera-based eye tracking labels served as training and testing labels for supplementary analyses (Fig. S5).

#### Dataset 5: Visual search

##### Data & task

These data were kindly provided by Julian and colleagues (Julian et al., 2018). Twenty-seven participants performed a self-paced visual search task, searching for the letter “L” in a search display filled with distractor letters “T”. Upon detection, participants pressed a button. Each trial lasted for an average of 7.50 seconds, followed by fixation at the screen center for 2 – 6 seconds. The number of distractors varied over trials between 81, 100, 144, 169, or 121. Participants performed either 4 or 6 runs of 6.5 minutes each. Task-relevant gaze positions were sampled within a window ofX= 17° and Y = 17° visual angle. Camera-based eye-tracking data were acquired and served as training and testing labels for our model (see below).

##### fMRI-data acquisition & preprocessing

Imaging data were acquired on a Siemens 3T MAGNETOM Prisma MRI scanner located at the Center for Functional Imaging in Philadelphia, USA. Following EPI-parameters were used: voxel size = 2mm isotropic, TR = 1000ms, TE = 25ms, flip angle = 45°, multiband factor = 4. Images were corrected for head motion using SPM12 and coregistered to our template space as described below. Note that the original dataset includes 9 more participants whose eyeballs were cut off on the functional images and that were excluded here.

##### Eye tracking

Gaze position was monitored at 30 Hz using the camera-based eye tracker LiveTrack AV by Cambridge Research Systems. We median-centered the time series and removed tracking noise by limiting the time series to values within the central 40 x 40 visual degree. We then split the data into individual scanning acquisitions to obtain the final gaze labels for model training and test.

#### Dataset 6: Fixation, smooth pursuit, free viewing & eyes-closed eye movements

##### Data & task

Four participants performed 4 viewing tasks while imaging data were acquired in the course of 9 scanning runs using 9 EPI-protocols (1 per run) along with concurrent camera-based eye tracking. First, they fixated sequentially at 37 locations on the screen for2s each starting in the screen center. The locations were determined using a custom random-walk algorithm that balanced the sampling of 12 directions (30 steps) and distances between the fixation points (4, 8, or 12 visual angle). Next, they performed a smooth pursuit version of this random-walk task for which we linearly interpolated the trajectory between fixation points. This resulted in a target moving sequentially into 12 directions at a speed of either 2/s, 4/s, or 6/s, changing to a randomly selected direction and speed every 2s. Next, participants freely explored 30 sequentially presented images of everyday objects for 3s each. The images were randomly drawn from the THINGS database (Hebart et al., 2019). Finally, participants closed their eyes and moved them either from left to right or from top to bottom fora total of 105 s. Switches between horizontal and vertical movements were indicated via button press.

##### fMRI-data acquisition & preprocessing

Imaging data were acquired using 9 EPI-sequences on a Siemens 3T MAGNETOM Skyra located at the St. Olavs Hospital in Trondheim, Norway. The sequences featured 3 repetition times and 3 voxel sizes in a 3×3 design. All images were corrected for head motion using SPM12 and coregistered to our template space as described below. See Table 2 for parameter details. Data acquisition was approved by the regional committees for medical and health research ethics (REC), Norway, and participants gave written informed consent prior to scanning.

**Table 2:** Sequence parameters of the 9 EPI-protocols used in the acquisition of dataset 6. For each sequence, we list the isotropic voxel size, the repetition time (TR), the echo time (TE), the flip angle (FA), the multiband factor (MB), partial Fourier factor (pF) and the number of slices (#Slices). All flip angles were aligned to the respective Ernst angle.

| Sequence | Voxel size | TR | TE | FA | MB | pF | #Slices |
| --- | --- | --- | --- | --- | --- | --- | --- |
| 1 | 1.5mm<sup>3</sup> | 1250ms | 26ms | 66 | 4 | 7/8 | 40 |
| 2 | 1.5mm<sup>3</sup> | 1800ms | 26ms | 74 | 4 | 7/8 | 40 |
| 3 | 1.5mm<sup>3</sup> | 2500ms | 26ms | 80 | 4 | 7/8 | 40 |
| 4 | 2.0mm<sup>3</sup> | 1250ms | 26ms | 66 | 4 | 7/8 | 60 |
| 5 | 2.0mm<sup>3</sup> | 1800ms | 26ms | 74 | 4 | 7/8 | 60 |
| 6 | 2.0mm<sup>3</sup> | 2500ms | 26ms | 80 | 4 | 7/8 | 60 |
| 7 | 2.5mm<sup>3</sup> | 1250ms | 26ms | 66 | 4 | 7/8 | 60 |
| 8 | 2.5mm<sup>3</sup> | 1800ms | 26ms | 74 | 4 | 7/8 | 60 |
| 9 | 2.5mm<sup>3</sup> | 2500ms | 26ms | 80 | 4 | 7/8 | 60 |

##### Eye tracking

Gaze position was monitored during the experiment at 1000 Hz using an Eyelink 1000 eye tracker. Tracking noise was reduced by excluding samples at which the pupil size diverged more than two standard deviations from the mean. Blinks were removed. The time series was downsampled to 60 Hz and median-centered based on the median gaze position of the free viewing condition within each scanning run. We then split the time series into individual scanning acquisitions to obtain the final training and testing gaze labels for our model.

### Eye masks, co-registration & normalization

Eye masks were created by manually segmenting the eyeballs including the adjacent optic nerve, fatty tissue and muscle area in the Colin27 structural MNI template using itkSNAP (http://www.itksnap.org, Fig. 1A). We then created a group-average functional template by averaging the co-registered functional images of 29 participants. These were acquired while the participants fixated at the screen center for around 13 minutes each in the course of a longer scanning session (Nau et al., 2018a). To ensure that the final eye masks contain the eyeballs of every participant, all imaging data underwent three co-registration steps conducted using Advanced Normalization Tools (ANTs) within Python (ANTsPy). First, we co-registered each participant’s mean-EPI non-linearly to our group-level average template. Second, we co-registered all voxels within a bounding box that included the eyes to a pre-selected bounding box in our group template to further improve the fit. Finally, we co-registered the eyeballs to the ones of the template specifically. Importantly, all data in our group-average template reflected gaze coordinates (0,0), i.e. the screen center. This third eyeball co-registration hence centered the average gaze position of each participant on the screen. We did this to improve the ft but also because it aligned the orientation of the eyeballs relative to the screen across participants. Finally, each voxel underwent two normalization steps. First, we subtracted the across-run median signal intensity from each voxel and sample and divided it by the median absolute deviation (MAD) over time (temporal normalization). Second, for each sample, we subtracted the mean across all voxels within the eye masks and divided by the standard deviation across voxels (spatial normalization). The fully co-registered and normalized voxels inside the eye masks served as model input.

### Model architecture

DeepMReye is a convolutional neural network that uses three-dimensional data to classify a two-dimensional output;the horizontal (X) and vertical (Y) gaze coordinates on the screen. The model uses the voxel intensities from the eye masks as input and passes it through a series of 3D-convolutional layers interleaved with group normalization and non-linear activation functions (mish, Misra 2019). In detail, the eye mask (input layer) is connected to a 3D convolutional block with a kernel size of 3 and strides of 1, followed by dropout and a 3D convolutional downsampling block which consists of one 3D-convolution followed by a 2×2×2 average pooling layer. After this layer, we use a total of six residuals blocks, in which the residual connection consists of one 3D convolutional block, concatenated via simple addition. Each residual block consists of group normalization, non-linear activation, and a 3D convolution, which is applied twice before being added to the residual connection. This results in a bottleneck layer consisting of 7680 units, which we resample to achieve sub-TR resolution (see details below). The time resolution dictates the number of resampled bottleneck layers, with e.g. 10 resampled layers producing a 10 times higher virtual resolution than the original TR. Each resampled layer is connected to a dense (fully-connected) layer which decodes the corresponding gaze position.

In addition to the above described model decoding gaze position directly, we also added a second block of fully-connected layers connected to the bottleneck layer. This second fully-connected layer block did not classify gaze position, but instead tried to predict the Euclidean error of the first model. This allowed us to obtain an unsupervised Euclidean error for each decoded gaze sample, even when test labels were missing. We refer to this predicted, unsupervised Euclidean error as the predicted error. It indicates how certain the model is about its own gaze decoding output and is strongly correlated with the real Euclidean error in our test data (Fig. 2B, Fig. S1). If the unsupervised error is high, the model itself anticipates that the decoded gaze position likely diverges much from the real gaze position. Accordingly, samples with high predicted error should not be trusted. DeepMReye is trained using a combination of the two losses, the Euclidean Error (90% weighting) and the predicted error loss (10% weighting) as described in detail below.

### Model optimization & training

Hyper-parameters were optimized using random search, which we monitored using the Weights & Biases’ model tracking tool (Biewald, 2020). Following parameters were optimized: the learning rate (0.001-0.00001), the number of residual blocks (depth, 3-6), the size of the filters (16-64), the filter multiplier per layer (1-2, e.g. 32, 64, 128 uses a multiplier of 2), the activation function (relu, elu, mish), the number of groups in the group normalization (4,8,16), the number of fully-connected layers (1,2), the number of units in each fully-connected layer (128-1024) as well as the dropout rate (0-0.5). In addition, to further improve the generalizability of our model, we added following data augmentations to the model training: input scaling, translations (X, Y, Z) and rotations (azimuth, pitch, and roll) which were applied on each sample.

We used Adam as learning algorithm (Kingma & Ba, 2015) and a batch size of 8 to train the model. Because considering samples from different participants improved model performance in an earlier version of our pipeline, we mixed samples in each training batch to represent 3D-inputs from different participants. For estimating the loss between real and predicted gaze position we used the Euclidean error:

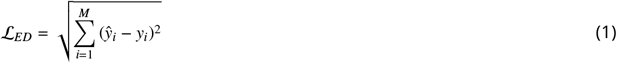

with *y_i_* the real gaze position, and *ŷ_i_*, the predicted gaze position. For calculating the predicted error, which reflects an unsupervised estimate of the Euclidean error, we used the mean squared error between real and predicted Euclidean error, which itself has been computed using the real and predicted gaze path as described above. The predicted error was computed as:

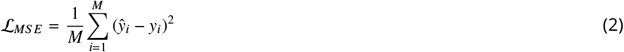

with *y_i_*, being the Euclidean error between real and predicted gaze path 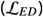, and *ŷ_i_*, being the predicted Euclidean error for this sample. The full loss for optimizing the model weights was computed as:

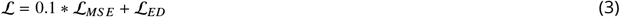

### Decoding schemes

We implemented three decoding schemes differing in how the data was split into training and test set. These decoding schemes are described in the following.

#### Within-participant decoding

Here, we split the data of each participant into two equally sized partitions (50/50% split). The model was trained on one half of the data of all participants and then tested on the other half of the data of all participants. This cross-validation procedure allowed the model to learn about the intricacies and details of each participant’s MR-signal and behaviors, while still having to generalize across participants and to new data of the same participants (Fig. S4).

#### Across-participant decoding

To test whether the model generalizes to held-out and hence fully independent participants, we further implemented an across-participant decoding scheme. This scheme represents our default pipeline and was used to obtain the main results Fig. 2. Each dataset was split into 5 equally sized partitions containing different participants. We then trained the model on 4 of these data partitions and then decoded from the fifth (80/20% split). This procedure was cross-validated until all data partitions and hence all participants were tested once. The across-participant decoding scheme requires the model to generalize to eyeballs and behavioral priors that it has not encountered during training. The fMRI and eye-tracking data however have been acquired on the same scanner and with the same scanning protocol.

#### Across-dataset decoding

Finally, we tested whether DeepMReye generalizes across datasets that have been acquired in independent participants performing different viewing tasks scanned on different scanners and with different scanning protocols. We trained the model in a leave-one-dataset-out fashion using all datasets (Fig. 2), meaning that the model was trained on all datasets except one and then tested on the one that was held out. This procedure was cross-validated until all datasets and hence all participants were tested once. Note that the voxel sizes and repetition times used for the acquisition of the key datasets 1-5 were similar, but that the model still had to generalize across different participants, MRI scanners and other scan parameters (e.g. slice orientation). Further note that the model performance oftheacross-dataset procedure would likely further improve if even more diverse viewing behaviors and fMRI data were used for model training (Fig. S4).

### Model quantification

To quantify model performance, we used the Euclidean error as described above for model training and evaluation. In addition, we computed the Pearson correlation and the *R*^2^-score as implemented in scikit-learn (Pedregosa et al., 2011) between real and decoded gaze path for model inference. The *R*^2^-score expresses the fraction-of-variance that our gaze decoding accounted for in the ground truth gaze path.

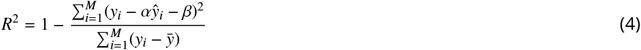

with *y_i_*, the ground truth of sample *i*, *ŷ_i_*, the predicted value and y the mean value. Unlike the Pearson correlation, or the squared Pearson correlation, the *R*^2^-score used here is affected by the scaling of the data and can be arbitrarily negative.

### Decoding from the eyeballs and early visual cortex with time-shifted data

To investigate if the decoding is instantaneous or further improves when temporal delays are being considered, we shifted the functional image time series relative to the gaze labels. We again used the free-viewing dataset (Julian et al., 2018), because it featured the most complex and natural viewing behaviors in our sample. For each image shift (0-10 TR’s), we retrained the full model and tested it on held-out participants using the across-participant decoding scheme.

To further assess whether DeepMReye can also be used to decode from brain activity directly, we used the same temporal shifting procedure while decoding from area V1. The regions-of-interest mask was obtained by thresholding the Juelich-atlas mask “Vi-sual_hOc1.nii” at 60 % probability and reslicing it to the resolution of our template space (Fig. S7). As the model is agnostic to the dimensions of the input data, decoding from region-of-interests other than the eyeballs required no change in model architecture.

### Effect of training set size

To evaluate how the number of participants in the training set influences decoding performance, we retrained the model using different subsets of participants across model iterations (1-21 participants). For each iteration, we tested the model on the same 6 participants, which were not part of the training set. To ensure that the results were robust and did not depend on individual details of single participants used for model training, we repeated this procedure 5 times for each training-set size and then averaged the results. To do so, we randomly assigned participants to the training set in each cross-validation loop while keeping the test set fixed. Moreover, to avoid overfitting to these small training sets, we reduced the number of training epochs, using *e* = 2 + *N* with *N* the number of participants in the current training run and e the number of epochs. We kept the number of gradient steps in each epoch constant (n=1500).

### Eyes-closed eye tracking

As a proof-of-concept, we tested whether DeepMReye is capable of decoding gaze position, or rather the state of the eyeballs, while the eyes are closed. We trained the model on the camera-based eye tracking labels of the 4 participants in dataset 6. We included the data acquired with all 9 scanning protocols and with all viewing behaviors tested (fixation, smooth pursuit, and picture viewing). We then evaluated the model on one participant, who was instructed to close the eyes and move them alternatingly from left to right or up and down. The participant indicated the direction of movement by pressing a button which was used to color the coordinates in (Fig. 3B). The participant performed this task nine times for one minute each. To reduce overfitting to the viewing behaviors in the training set, we here used a higher dropout rate in the fully connected layers (drop ratio=0.5) than in our default model (drop ratio=0.1).

### Decoding at sub-imaging temporal resolution

Because different imaging slices are being acquired at different times, and because the MR-signal of a voxel could be affected by eye motion within each TR, we tested whether our model is capable of decoding gaze position at sub-imaging temporal resolution. Across different model iterations, we re-trained and re-tested the model using different numbers of gaze labels perTR (n = 1-10 labels), each time testing how much variance the decoded gaze path explained of the true gaze path. Decoding different numbers of gaze labels perTR was achieved by replicating the bottleneck layer n times, each one decoding gaze position for their respective time points using a fully connected layer. Importantly, the weights between these layers were not shared, which allowed each layer to utilize a different node in the bottleneck layer. Each layer could therefore capture unique information at its corresponding within-TR time point to decode its respective gaze label. To keep the overall explainable variance in the test set gaze path constant, we always upsampled the decoded gaze path to a resolution of 10 labels per TR using linear interpolation before computing the *R*^2^-score scores for each model iteration. Potential differences in model performance across iterations can therefore not be explained by differences in explainable variance. These final test *R*^2^-scores were range-normalized within each participant for visualization (Fig. 2F).

### Functional imaging analyses

We tested whether the decoding output of DeepMReye is suitable for the analysis of functional imaging data by regressing it against brain activity using a mass-univariate general linear model (GLM). This analysis was expected to uncover brain activity related to eye movements in visual, motion, and oculomotor regions. To demonstrate that our approach is applicable even for natural and complex viewing behavior, we conducted these analyses on the visual search dataset (Julian et al., 2018).

First, we decoded the median gaze position at each imaging volume using all cross-validation schemes described above. We then obtained an approximate measure of eye movement amplitude by computing the vector between gaze positions of subsequent volumes. Based on the vector length, or the amplitude of decoded putative eye movements, we built two regressors of interest;one for far eye movements (>66th percentile of movement amplitudes) and one for short eye movements (<33rd percentile of amplitudes). The mid-section was excluded to separate the modeled events in time. The two resulting regressors per scanning run were binarized and convolved with the hemodynamic response function implemented in SPM12 using default settings. Head-motion parameters obtained during preprocessing were added as nuisance regressors. Contrasting the resulting model weight between far and short eye movements yielded one t-statistics map per participant.

To test which brain areas signaled the difference between far and short eye movements, we normalized the t-map of each participant to MNI-space and smoothed it with an isotropic Gaussian kernel of 6mm (full-width-half-maximum). The smoothed statistical maps were then used to compute an F-statistic on group level using SPM12. Moreover, to compare the results obtained with DeepMReye to the ones of conventional eye tracking we repeated the imaging analysis described above using gaze positions obtained with a conventional camera-based eye tracker. The final F-statistics maps were warped onto the fsaverage Freesurfer template surface for visualization using Freesurfer (https://surfer.nmr.mgh.harvard.edu/).

## Supplementary Material

**Figure S1:**
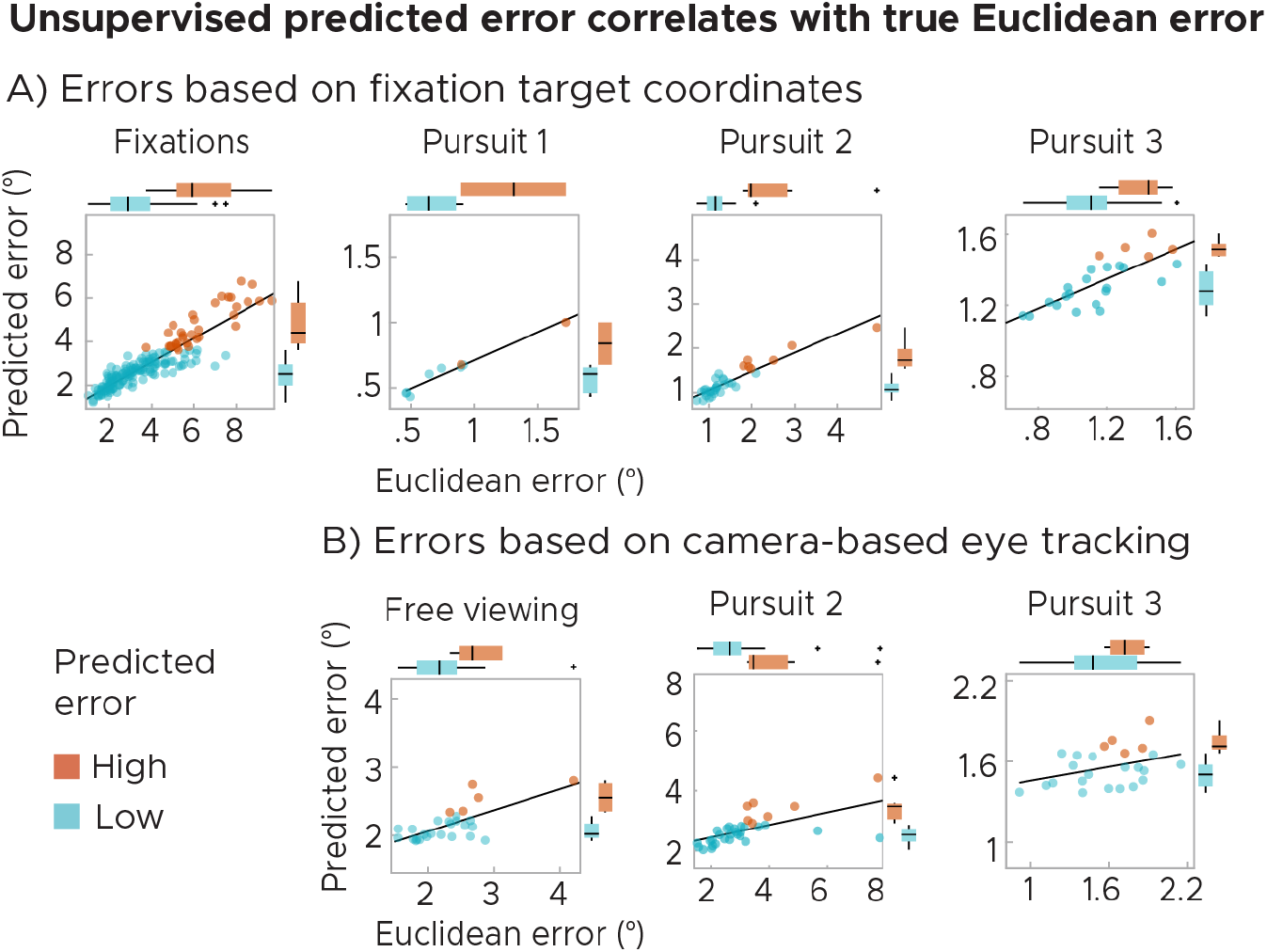
Predicted error (PE) correlates with the Euclidean error between real and predicted gaze positions. This allows to filter the test set post-decoding based on estimated reliability. A) Results plotted for models trained and tested using the fixation target coordinates. B) Results plotted for models trained and tested using labels acquired using camera-based eye tracking. We plot single-participant data with regression line. Participants were split into 80% most reliable (Low PE, blue) and 20% least reliable participants (high PE, orange). All scores expressed in visual degrees.

**Figure S2:**
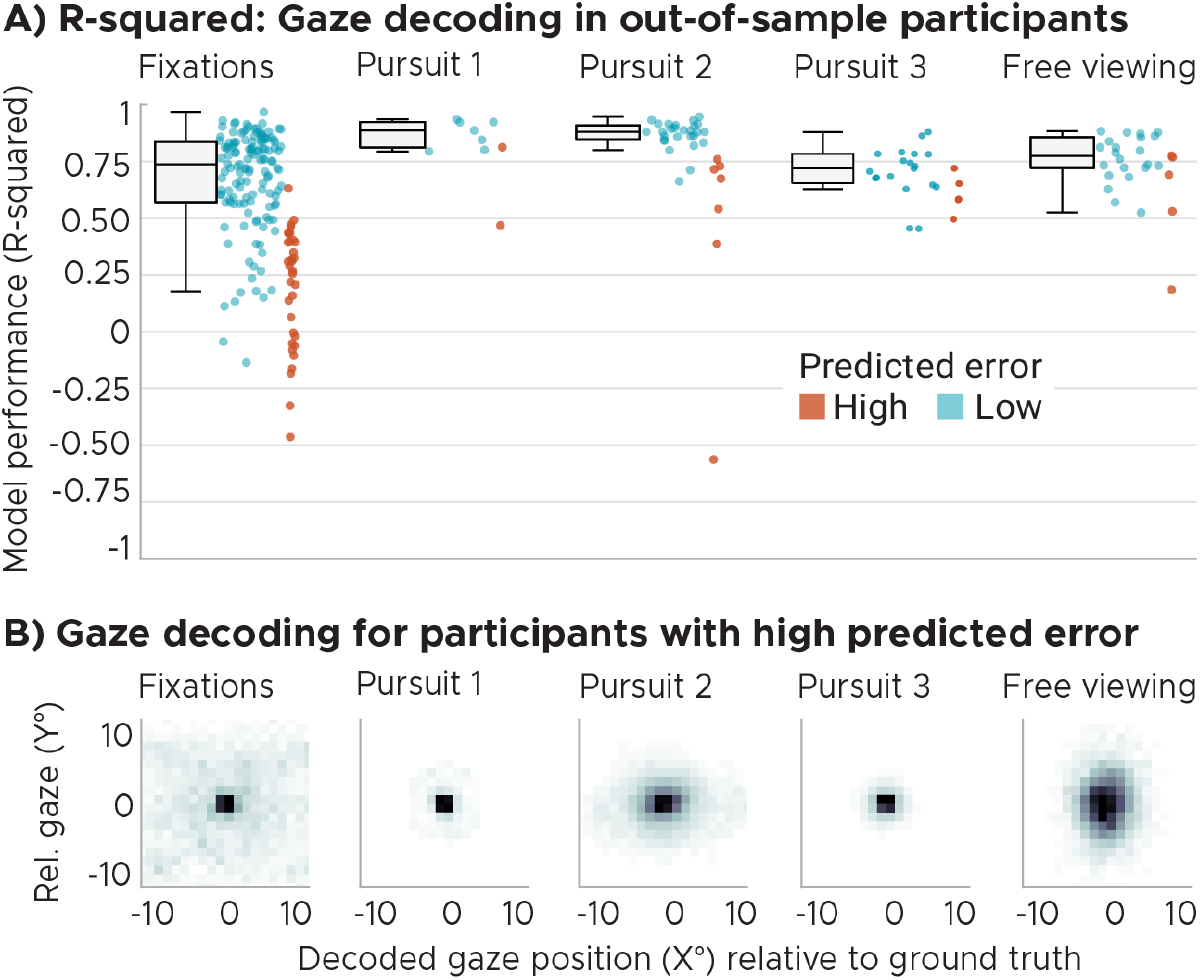
A) Gaze decoding group results expressed as the coefficient-of-determination (*R*^2^). Top panel shows gaze decoding expressed as the *R*^2^-score implemented in scikit-learn (Pedregosa et al., 2011) between the true and decoded gaze trajectory for the five key datasets featuring fixations, 3x smooth pursuit and visual search. Note that *R*^2^ can range from negative infinity to one. Participants are color coded according to predicted error (PE). We plot Whisker-box-plots for Low-PE participants and single-participant data for all. (B) Group-average spread of decoded positions around true positions collapsed over time in visual degrees for participants with high predicted error (orange dots in A).

**Figure S3:**
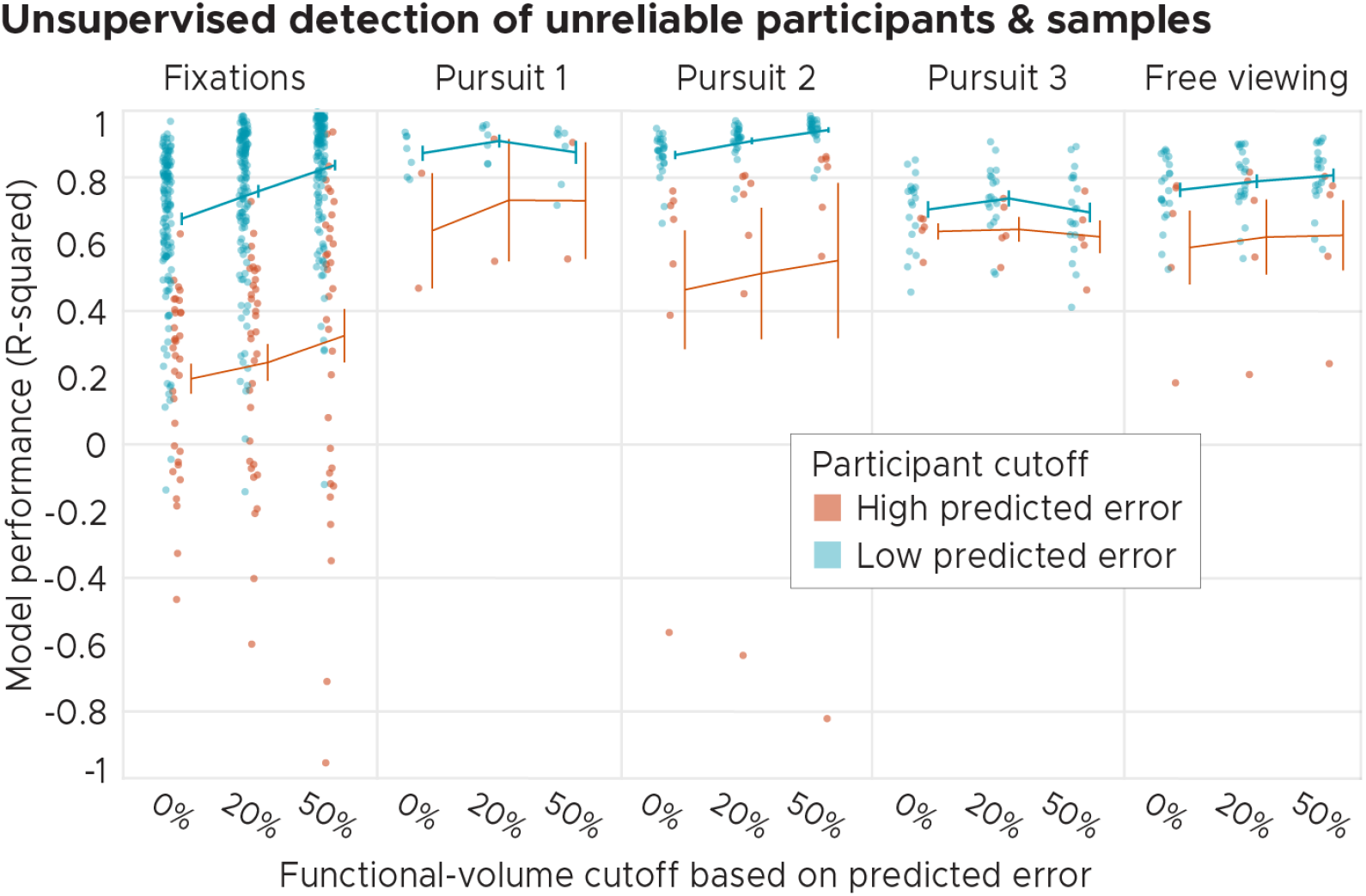
Model performance evaluated before and after exclusion of volumes with unreliable decoding. Here, before computing model performance we filtered out either the 0%, 20% or 50% least reliable volumes (i.e. those with the highest predicted error (PE)). Model performance is expressed as the coefficient-of-determination *R*^2^-score implemented in scikit-learn (Pedregosa et al., 2011) between true and decoded gaze trajectory for the five key datasets featuring fixations, 3x smooth pursuit and visual search. Note that *R*^2^ can range from negative infinity to one. We plot single participant data (dots) as well as the mean ± standard error of the mean. Participant dots were additionally color coded according to the participants’ PE.

**Figure S4:**
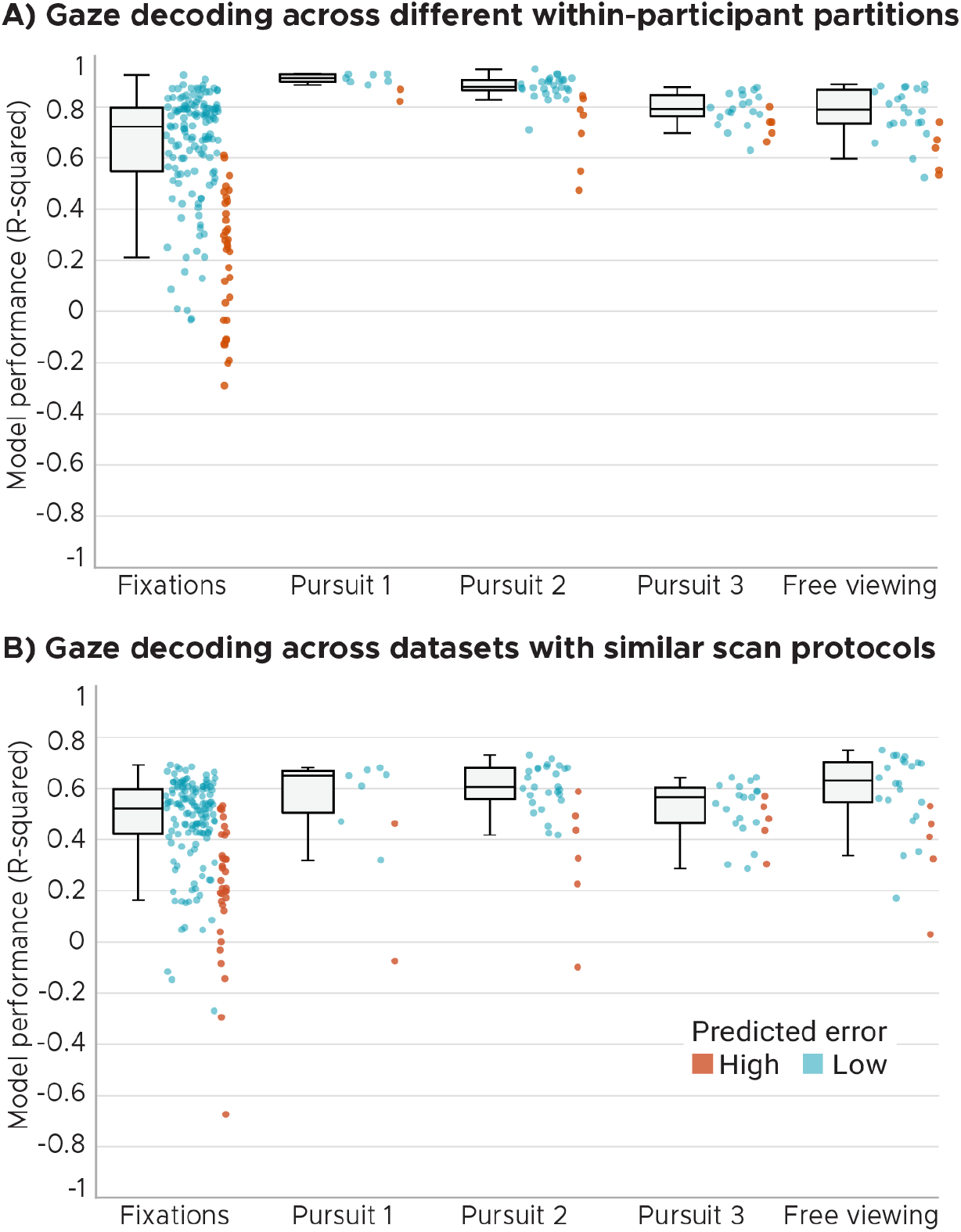
A) Within-participant gaze decoding obtained by training and testing the model on different data partitions of all participants within a dataset. B) Across-dataset gaze decoding obtained using leave-one-data-set-out cross-validation. We plot the *R*^2^-score as implemented in scikit-learn (Pedregosa et al., 2011) between true and decoded gaze trajectory for the five key datasets featuring fixations, 3x smooth pursuit and visual search. Note that *R*^2^ can range from negative infinity to one. The results of datasets 1-3 were obtained using the fixation target labels, the ones of datasets 4-5 were obtained using camera-based eye tracking labels. Participants are color coded according to predicted error (PE). We plot Whisker-box-plots for Low-PE participants and single-participant data for all.

**Figure S5:**
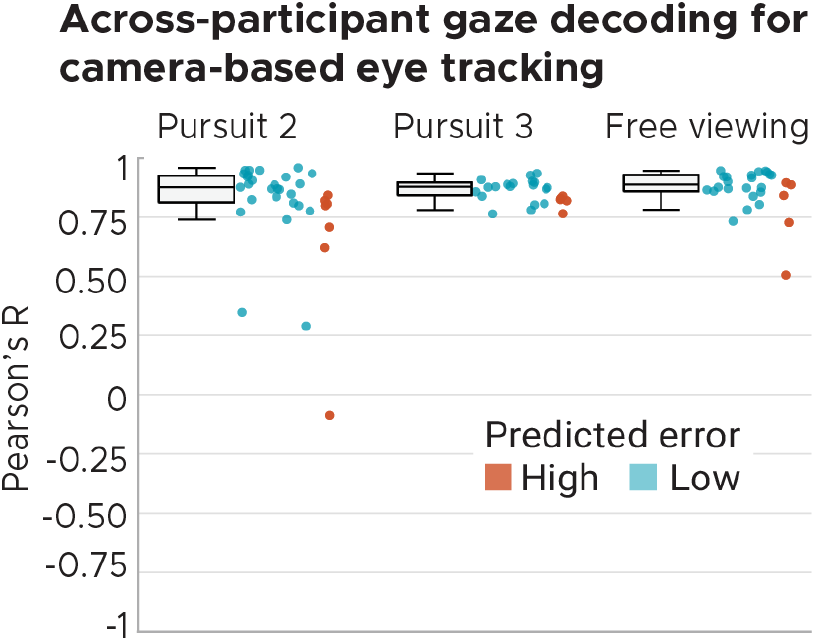
Gaze decoding evaluated using camera-based eye tracking for smooth pursuit datasets 3-4. Model performance expressed as the Pearson correlation between true and decoded gaze trajectory for the datasets with camera-based eye tracking. Because the visual search dataset 5 used labels obtained using camera-based eye tracking as well, we additionally plot the results obtained for this dataset again for the sake of completeness. Participants are color coded according to predicted error (PE). We plot Whisker-box-plots for Low-PE participants and single-participant data for all.

**Figure S6:**
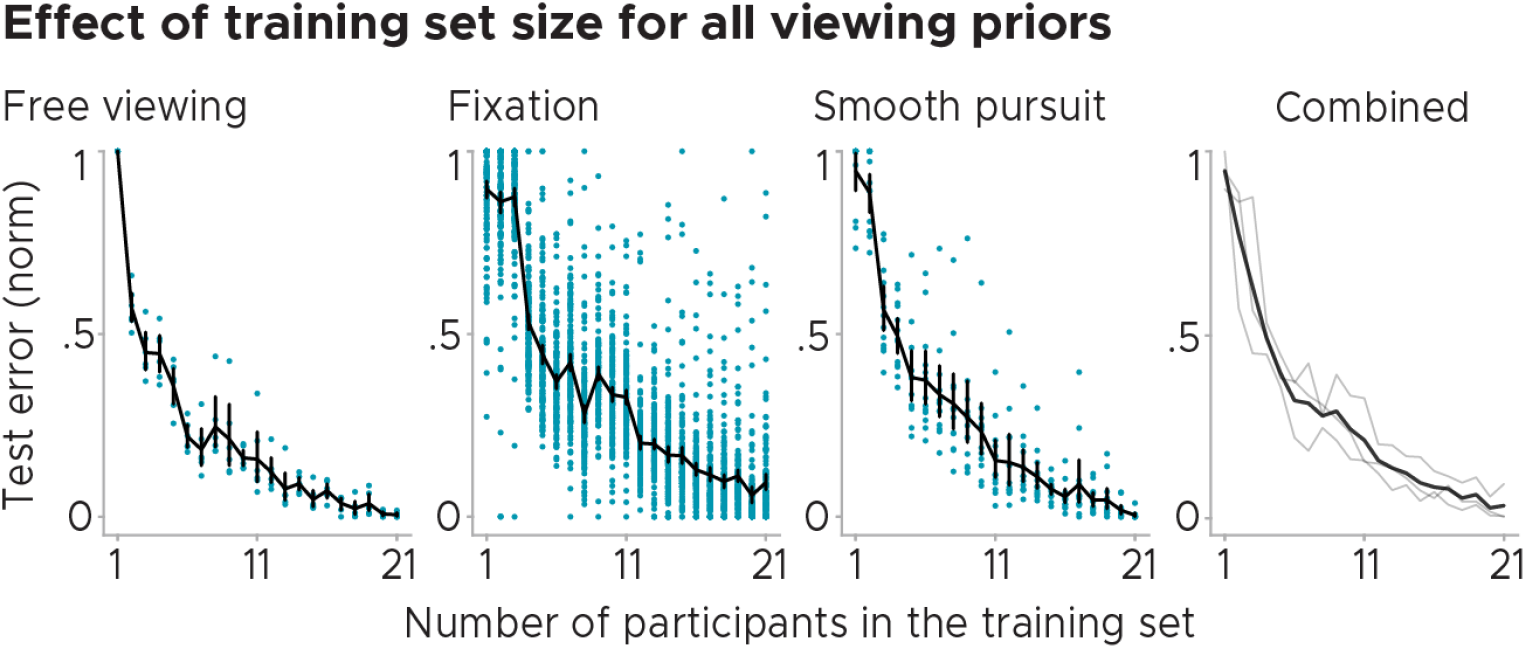
Normalized test error as a function of how many participants were used for model training plotted for three different viewing behaviors. We plot single participant data (dots) as well as the across-participant average model performance (black lines). Error bars depict the standard error of the mean. Right panel shows the average across datasets.

**Figure S7:**
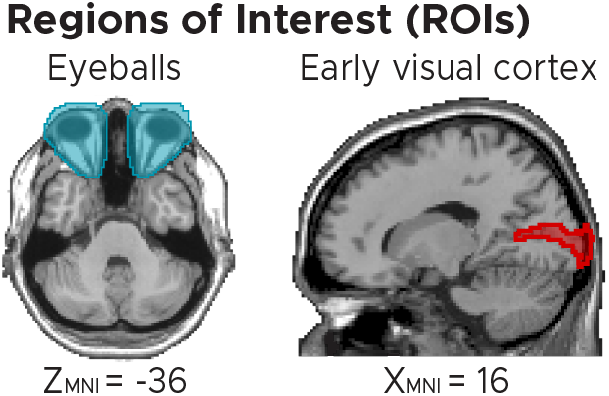
Visualisation of eyeball and visual cortex (V1) masks used for decoding in Figure 2E. Eyeballs were manually segmented in the structural scan of the SPM-template participant “Colin27”. The V1 mask was obtained by thresholding the Juelich-atlas mask “Visual_hOc1.nii” at 60 percent probability. MNI coordinates added. For decoding, both masks were resliced to 2mm isotropic to match the voxel resolution of our template space.

**Figure S8:**
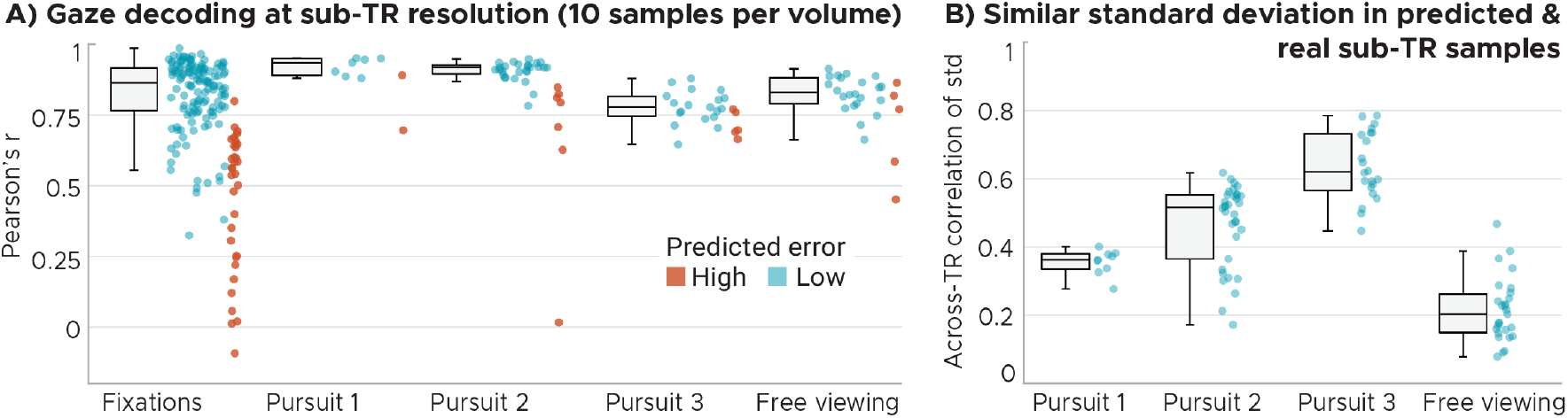
Sub-imaging decoding resolution. A) Group results when all 10 sub-TR samples are considered for computing the Pearson correlation between true and decoded gaze trajectories. Participants are color coded according to predicted error (PE). We plot Whisker-box-plots for Low-PE participants and single-participant data for all. B) Similar standard deviation of real and decoded gaze labels within each functional volume (TR), i.e. if the 10 real gaze labels of a TR had a high standard deviation (indicating larger eye movements within this TR) then the 10 decoded gaze labels showed a high standard deviation as well. We plot the Pearson correlation between the within-TR standard deviation computed using the full time course of each participant as Whisker-box-plots and single-participant data as dots.

**Figure S9:**
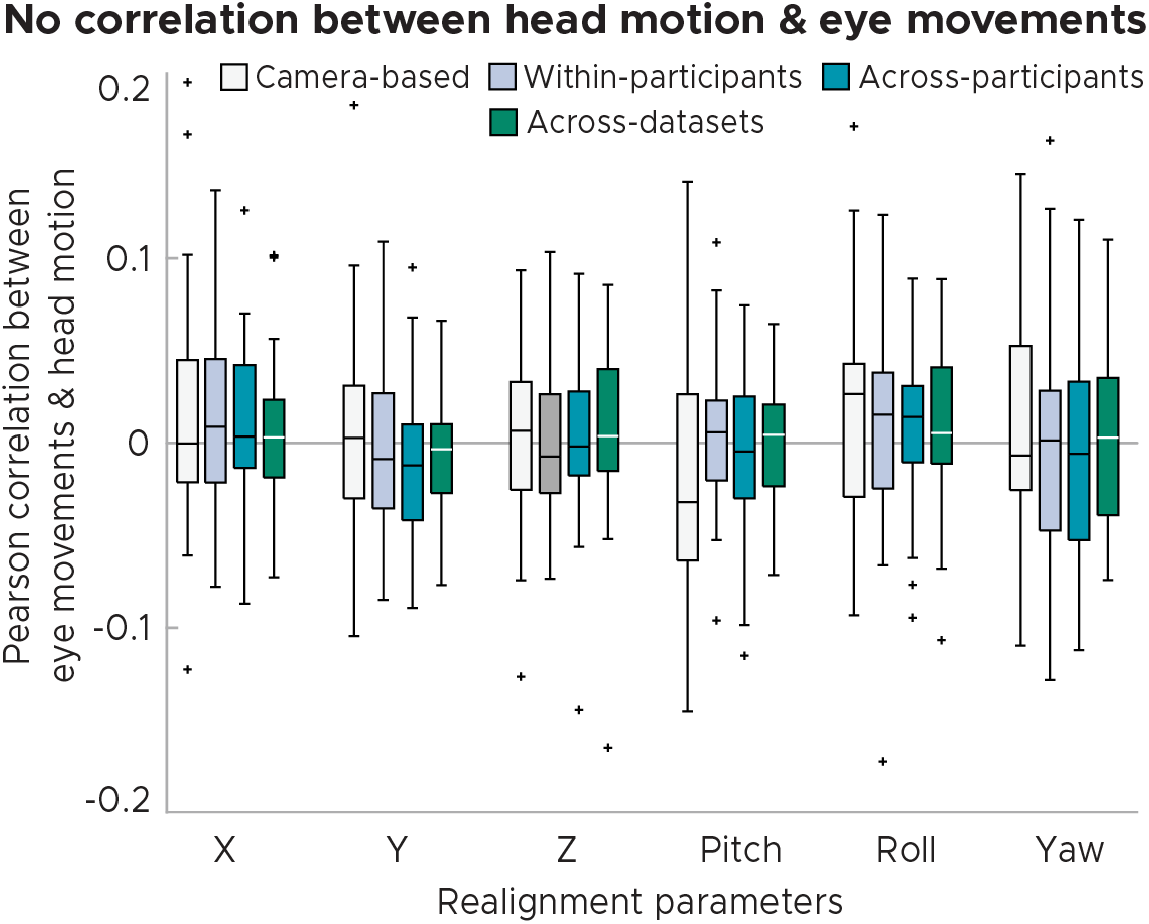
No correlation between eye movements and head motion in visual search dataset 5. Eye movements were computed as the vector length between gaze positions of subsequent volumes. Head motion estimates reflect the 6 SPM12-realignment parameters. We plot Whisker-box-plots of this correlation computed for gaze labels obtained with camera-based eye tracking as well as with three cross-validation schemes of DeepMReye (within-participant-, across-participant- and across-dataset prediction).

**Figure S10:**
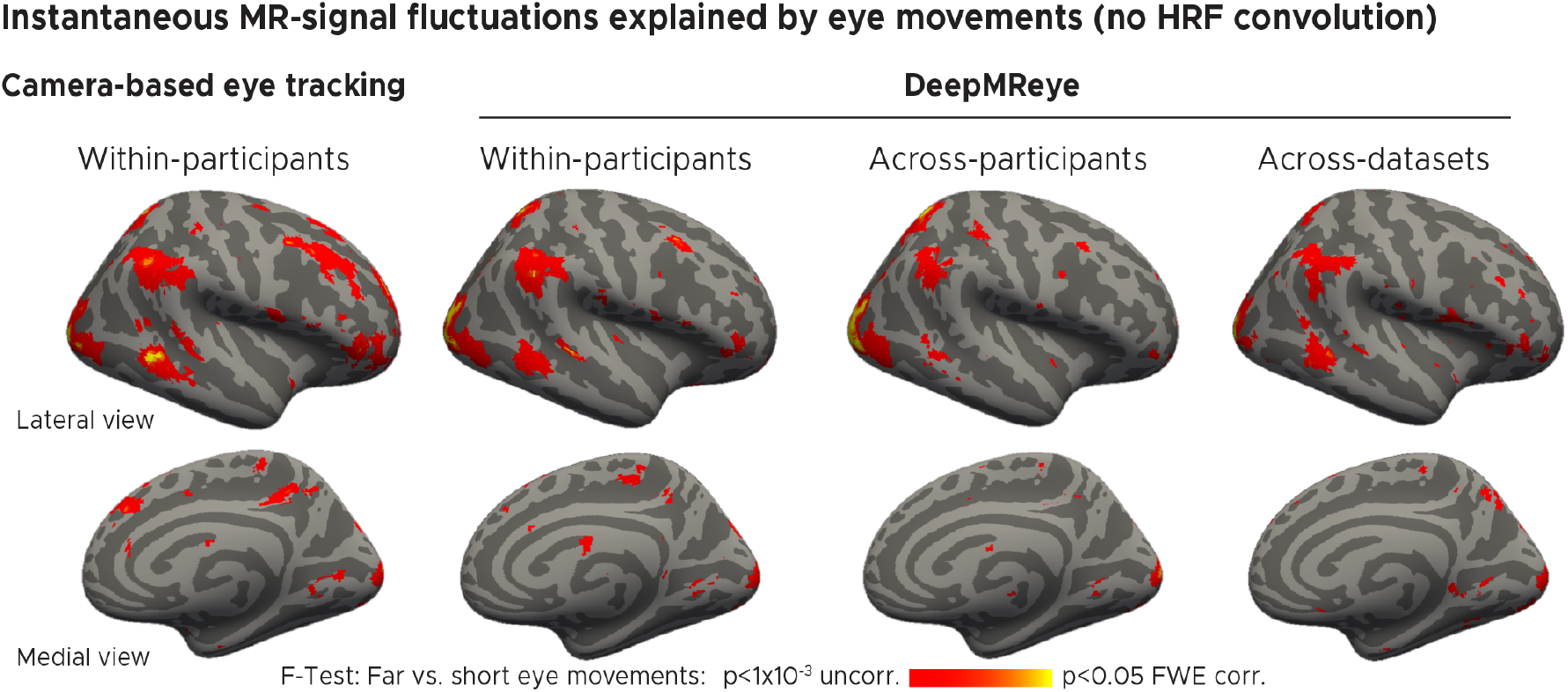
General-linear-model (GLM) group results for the contrast’Far vs. short eye movements’during visual search without accounting for the hemodynamic response function. We plot the F-statistic of this contrast superimposed on a template surface (fsaverage) for gaze-labels obtained with camera-based eye tracking (first panel) as well as for three DeepMReye cross-validation schemes. Within-participants: All participants of a dataset were included with different partitions in model training and test. Across-participants: Different participants were included during model training and test. Across-datasets: Different datasets (and hence also different participants) were included during model training and test.

